# Astrocytic Ryk signaling coordinates scarring and wound healing after spinal cord injury

**DOI:** 10.1101/2024.10.16.618727

**Authors:** Zhe Shen, Bo Feng, Wei Ling Lim, Timothy Woo, Yanlin Liu, Silvia Vicenzi, Jingyi Wang, Brian K. Kwon, Yimin Zou

## Abstract

Wound healing after spinal cord injury involves highly coordinated interactions among multiple cell types, which is poorly understood. Astrocytes play a central role in creating a border against the non-neural lesion core. To do so, astrocytes undergo dramatic morphological changes by first thickening the processes and then elongating and overlap them. We show here show that the expression of a cell-surface receptor, Ryk, is induced in astrocytes after injury in both rodent and human spinal cord. Astrocyte-specific knockout of Ryk dramatically elongated the reactive astrocytes and accelerated the formation of the border and reduced the size of the scar. Astrocyte-specific knockout of Ryk also accelerated the injury responses of multiple cell types, including the resolution of neuroinflammation. Single cell transcriptomics analyses revealed a broad range of changes cell signaling among astrocytes, microglia, fibroblasts, endothelial cell, etc, after astrocyte-specific Ryk knockout, suggesting that Ryk not only regulates the injury response of astrocytes but may also regulate signals which coordinate the responses of multiple cell types. The elongation is mediated by NrCAM, a cell adhesion molecule induced by astrocyte-specific conditional knockout of Ryk after spinal cord injury. Our findings suggest a promising therapeutic target to accelerate wound healing and promote neuronal survival and enhance functional recovery.

---

Traumatic spinal cord injury starts with primary injury to neurons and glia and continues with a much larger secondary injury caused by inflammation, ischemia, excitotoxicity and oxidative stress, which ends with the formation of a fibrotic scar bordered by astrocytes (1). Injured axons initially retract from the lesion site due to the repulsive function of the reinduced guidance cues, such as the Wnt family proteins, and cannot grow back through the fibrotic scar (2–5). Injured axons and their collateral branches usually show significant growth around the injury site or elsewhere away from the injury site and sometimes the axons can extend beyond the injury site through the spared tissue bypassing the lesion site (4, 5). Therefore, increasing the protection of the spared spinal cord tissue and promoting the regenerative growth of axons can enhance the repair of circuits and recovery of sensorimotor functions after spinal cord injury. The understanding of the role of astrocytes after spinal cord injury has been evolving and more recent studies highlight the benefit of the astrocytes in forming a protective border (6, 7). Reactive astrocytes undergo extensive morphological changes after injury, including initial thickening and then polarization/elongation and finally overlapping of the processes to line up the border (8).

The signaling mechanisms regulating these morphological changes are not known. In addition, astrocytes have complex roles in the inflammatory responses, both proinflammatory and anti-inflammatory, likely temporally and spatially controlled. How these processes are controlled in the astrocytes are not well understood (9). Finally, astrocytes are known to communicate with many other cell types through extensive cell-cell signaling (9). What are these signals and how do these communications affect wound healing are not clear. Elegant work lead to the discovery of the key signaling axis, which initially detect the injury and tissue damage and triggers astrogliosis, the TLR4/GP130/STAT3 axis (8). How STAT3 activates the downstream signaling mechanisms to mediate all the aforementioned events is unknown. Here, we report that Ryk, a receptor in both canonical and non-canonical Wnt signaling pathways, a direct transcriptional target of STAT3, plays central roles in regulating astrocyte morphology, differentiation and communication with multiple cell types for wound healing after spinal cord injury.

## RESULTS

### Ryk expression is induced in injured spinal cords in mice and humans

A Wnt receptor, Ryk, known to mediate axon repulsion in development as well as in injured adult spinal cord, was also observed induced in the lesion area outside axons, particularly on the border of the developing scar, after spinal cord injury (2, 5, 10). We therefore analyzed Ryk expression and asked whether it is expressed in reactive or border-forming astrocytes. To validate Ryk staining and astrocyte identity, we crossed *Ryk* floxed allele with *hGFAPcre^ERT2^* or *Aldh1L1 cre^ERT2^* and performed C5 dorsal column lesion (5). Mice were subjected to cervical 5 (C5) dorsal column lesion and co-stained with a polyclonal Ryk antibody we previously made in the lab and an astrocyte marker, GFAP, or a fibroblast marker, Fibronectin, at 1, 3,7 and 14 days after injury (Fig. 1A) (2, 11). Studies reported that astrocyte density was significantly increased within 250 µm from the lesion border (8, 12). Therefore, we examined Ryk expression in those areas and we found that Ryk significantly was increased at 1, 3 and 7 days after spinal cord injury and started to decrease at 14 days after injury (Fig. 1B). And the Ryk immunoreactivity was found co-localized with astrocyte marker, GFAP (Fig. 1B, shown in the blue boxes).

**Fig. 1.**
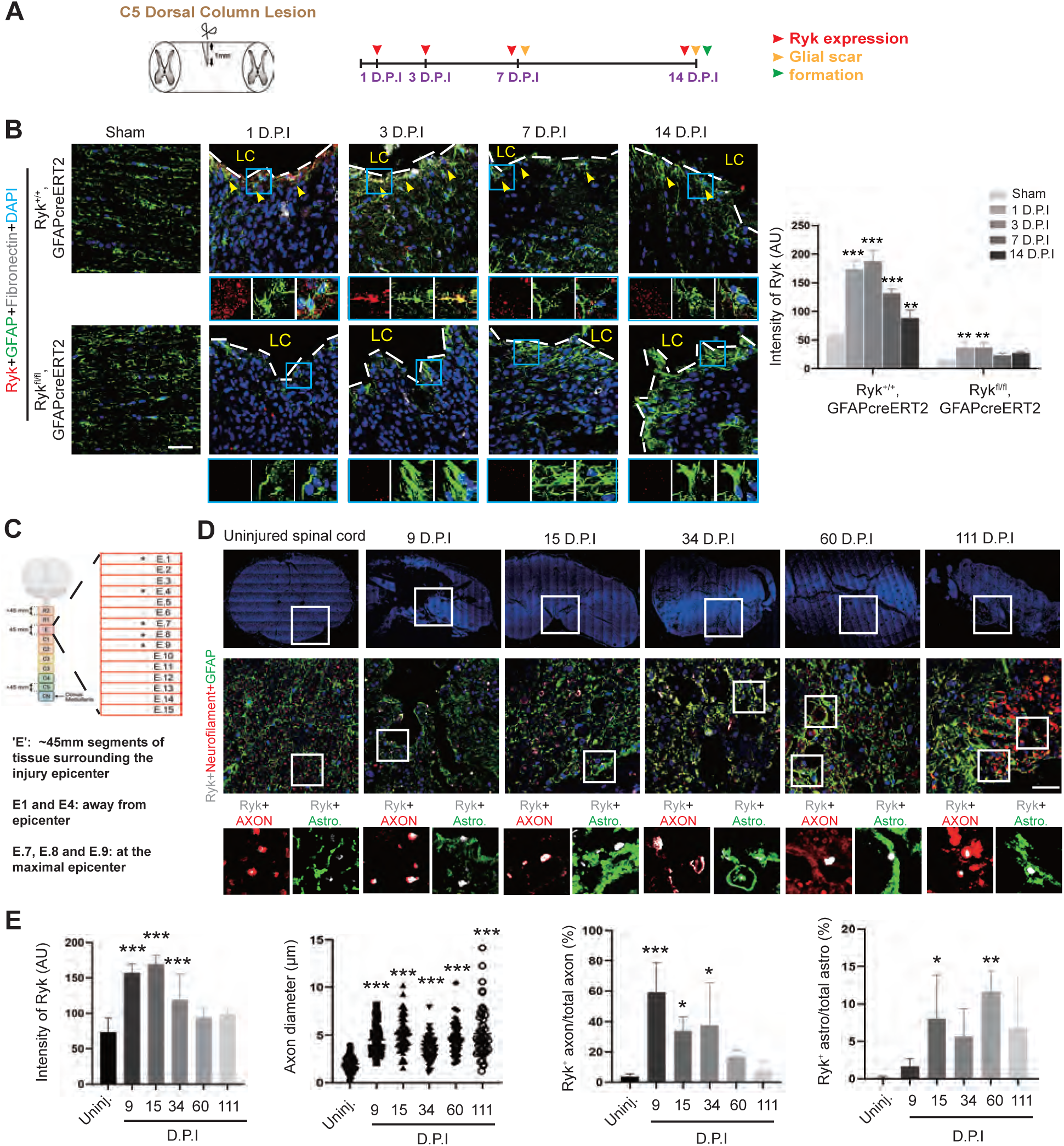
Induction of Ryk expression at injury sites in mice and humans after spinal cord injury. **(A)** Experimental design for mice spinal cord injury. **(B)** Immunohistochemistry with antibodies against Ryk, GFAP and Fibronectin 1,3, 7 or 14 days after C5 dorsal column lesion in control or in *Ryk^fl/fl^* crossed with *GFAPcre^ERT2^*. N=3 for each group. Scale bar =40 µm. **(C)** Schematic showing the location of the human spinal cord tissue sections relative to the lesion epicenter. **(D)** Immunohistochemistry at different time points after human spinal cord injury with antibodies against Ryk (Grey), Neurofilament H (Red) and GFAP (Green). **(E)** Bar graphs showed quantitative analyses of Ryk expression levels, axon diameter, ratio of Ryk^+^ axons to total axons and ratio of Ryk^+^ astrocytes to total astrocytes. Scale bar =40 µm. Data are expressed as mean ± SD. \**P* < 0.05, \*\**P* < 0.01 and \*\*\**P* < 0.001 vs. the indicated groups.

Compared with WT animals, we found that Ryk expression was indeed abolished in astrocytes in *cKO* mice (Fig. 1B) from 1 to 14 days after injury, confirming the specificity of the Ryk antibody and the effectiveness of the Ryk *cKO* in astrocytes.

To test the relevance of Ryk function in human spinal cord injury, we analyzed spinal cord samples from 5 different patients at different time points of injury. From each patient, we analyzed 5 coronal sections (Fig. S1A). These sections were taken from Sections E1, 4, 7, 8 and 9. E7, 8 and 9 are the epicenter of the injuries (Fig. 1C). E1 is about 15 mm away from the lesion center while E4 is about 6 mm away from the core. First, we co-stained with a monoclonal Ryk antibody with an astrocyte marker GFAP and an axon marker Neurofilament in the lesion core (Sections E7, 8, 9) and found that Ryk was significantly increased in spinal tissues at 9 D.P.I. (C6/7 fracture dislocation/C6 AIS A),15 D.P.I. (C3/4 hyperextension injury/C4 AIS C) and 34 C.P.I (C6/7 fracture dislocation/C7 AIS A) in both astrocytes and in axons compared with spinal cord tissue from uninjured control, and the signal started to decrease at 60 D.P.I. (C4/5 hyperextension injury C4 AIS D) and 111 D.P.I. (C4/5 hyperextension injury C5 AIS A) (Fig. 1D and 1E). The average diameter of Neurofilament signal was found increased in the lesion core, suggesting axonal swelling after injury. Therefore, Ryk is induced on axon at the lesion core shortly after injury and the expression persists for at least 1 month but declines 2 months after injury (Fig. 1D and 1E). The expression of Ryk on the astrocytes, on the other hand, continued to increase but peaked at 2 months after injury. We then investigated the Ryk expression away from the lesion core with astrocyte marker SOX9 (in E4) (Fig. S2A) and axon marker Neurofilament (in E1) (Fig. S2B). We found that Ryk expression was significantly increased at 9, 15 and 34 days after injury and started to decline at 60 and 111 days in both E1 and E4 sections (Fig. S2C). And in E4 and E1 sections, Ryk immunoreactivity co-localized with the astrocyte marker SOX9 and the axonal marker Neurofilament, respectively.

### Astrocyte-specific *Ryk* conditional knockout resulted in smaller lesion volume, more preserved neurons and axons and sensory-motor functions

To explore role of the induced Ryk in astrocytes, we performed a number of analyses in the injured spinal cords of the astrocyte-specific *Ryk* conditional knockout mice. First, we measured the injury lesion volume based on GFAP delineation and observed a significant decrease in lesion volume 14 days after injury in astrocyte-specific *Ryk* knockouts using either *hGFAP-CreERT2* or *Aldh1l1-CreERT2* (Fig. 2A-2B). We then asked with the smaller lesion volume promotes the survival of neural tissues and observed that the neuronal marker NeuN was significantly increased 7 and 14 days after injury (Fig. 2C-2E), indicating astrocyte-specific *Ryk cKO* may have promoted neuronal survival. Since *Ryk cKO* reduced the lesion size, we further investigated whether there is any long-term functional benefit. Using grip strength and rotarod performance, we found that *Ryk cKO* protected the motor function for 2 months after C5 dorsal column lesion injury (Fig. 2F and 2G, n=6 for control group and n=7 for Ryk cKO). And we further analyzed the axon morphology 2 months after injury by injecting BDA into the motor cortex and co-stained GFAP and fibronectin. Using a digital stereotactic injector (Item: 51709, stoelting Co. USA), 0.5 µl of biotin dextran amine (BDA; MW 10,000; 10% in PBS; Molecular Probes) was injected into one of the 10 total sites (5 sites/side). The coordinates are 1.5 mm lateral to the bregma (mediolateral), 1.0, 0.5, 0, 0.75 and 1.5 mm from the bregma (anteroposterior) and 0.5 mm from the cortical surface (dorsoventral). We found that *Ryk cKO* significantly reduced the corticospinal tract axon retraction (Fig. 2H, normalized to pyramidal labeling) and increased the axon branch (Fig. 2I and 2J). Therefore, astrocyte-specific *Ryk cKO* is beneficial.

**Fig. 2.**
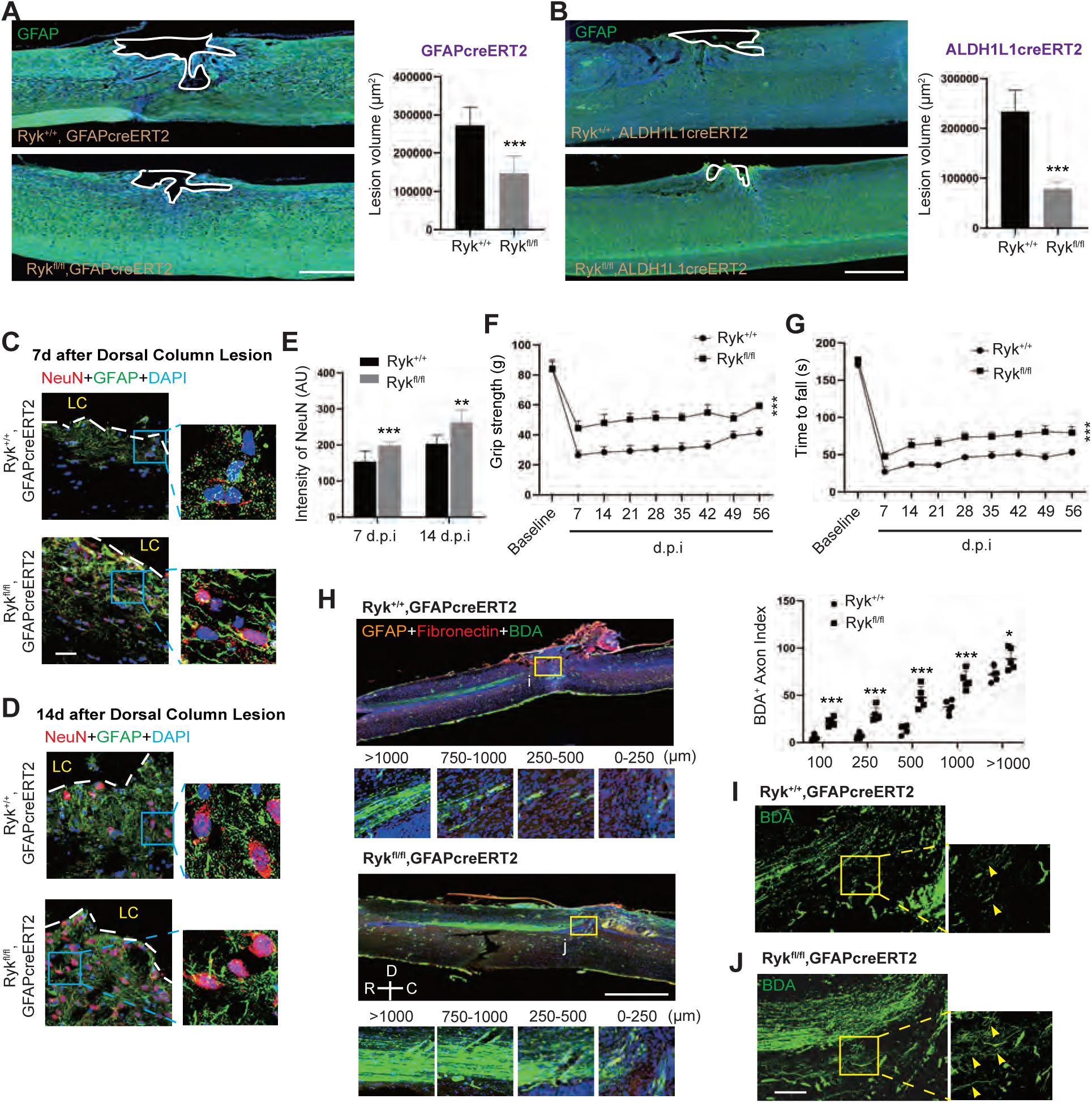
Smaller lesion volume, more preserved neurons and axons and sensory-motor functions in astrocyte-specific Ryk conditional knockout . **(A)**Lesion volume shown in sagittal sections of spinal cord stained with GFAP antibody 14 days after SCI in control or *Ryk^fl/fl^*crossed with *GFAPcre^ERT2^*. Scale bar =500 µm. N=5 for each group. **(B)** Lesion volume shown in sagittal sections of spinal cord stained with GFAP antibody 14 days after SCI in control or *Ryk^fl/fl^* crossed with *Aldh1L1cre^ERT2^*. Scale bar =500 µm. N=5 for each group. Immunohistochemistry with NeuN in spinal cord sections 7 days **(C)** or 14 days **(D)** after SCI stained. Scale bar =40 µm. **(E)**Quantification of **C** and **D**. Measurements of grip strength **(F)** and Rotarod performance **(G)** 2 months after C5 dorsal column lesion. N=6 for control group and n=7 for Ryk cKO. **(H)** Representative images of BDA-labeled CST axons with GFAP and fibronectin staining in control or *Ryk cKO* mice. Scale bar =1000 µm. N=5 for each group. Higher magnification of images for control **(I)** or *Ryk cKO* (J) from boxed regions (750 µm rostral to the lesion site) indicated in **(H).** Scale bar =50 µm. Data are expressed as mean ± SD. \**P* < 0.05, \*\**P* < 0.01 and \*\*\**P* < 0.001 vs. the indicated groups.

### Astrocyte-specific *Ryk* conditional knockout promoted polarization and elongation of astrocyte processes and accelerated astrocyte border formation

To better understand the role of Ryk in astrocytes and how astrocyte-specific Ryk knockout affect injury responses, we further analyzed the astrocyte border. In addition to the smaller lesion size, we found that the astrocytes form astrocyte border more efficiently in astrocyte-specific *Ryk cKOs* 14 days after injury (Fig. 3A), suggesting that astrocyte-specific *Ryk cKO* accelerated the injury response of astrocytes. In support of this, we found astrocyte processes were greatly elongated and highly polarized towards the injury site in *Ryk cKO*, particularly obvious 7 days after SCI when the density of GFAP-positive reactive astrocytes are still sparse and when it is possible to clearly observe their morphology (Fig. 3B and S3, mainly quantified in areas with sparse GFAP staining). These highly polarized astrocyte cell processes were also more overlapped in *Ryk cKO* (Fig. 3B and S3). Overlapping of processes is one of the key morphological features of border-forming astrocyte (BF). To further test these, we tested an astrocyte marker, SOX9, and found that SOX9 level was indeed significantly increased at 7 days after SCI in *Ryk cKO* (Fig. S4A and S4B), indicating *Ryk cKO* accelerated the process of reactivated astrocyte transforming to border-forming astrocytes. Furthermore, to test whether the change of astrocyte border in *Ryk cKO* caused by astrocyte proliferation, mice were administered a single injection of BrdU on each day from 2 to 7 d after SCI. And we found that the newly proliferated astrocyte significantly increased at 7 days after injury (Fig. S4C). However, there was no significant difference between WT and Ryk cKO in the number of proliferated astrocytes at 14 days after injury, suggesting that we are observing an acceleration of proliferation rather than an overall increase (Fig. S4D). These data suggested that Ryk cKO accelerated the proliferation of reactive astrocytes after injury to form the border, which was observed at Day 7. Once enough astrocytes were generated, proliferation slowed down and the final number of astrocytes is similar to wildtype control at Day 14.

**Fig. 3.**
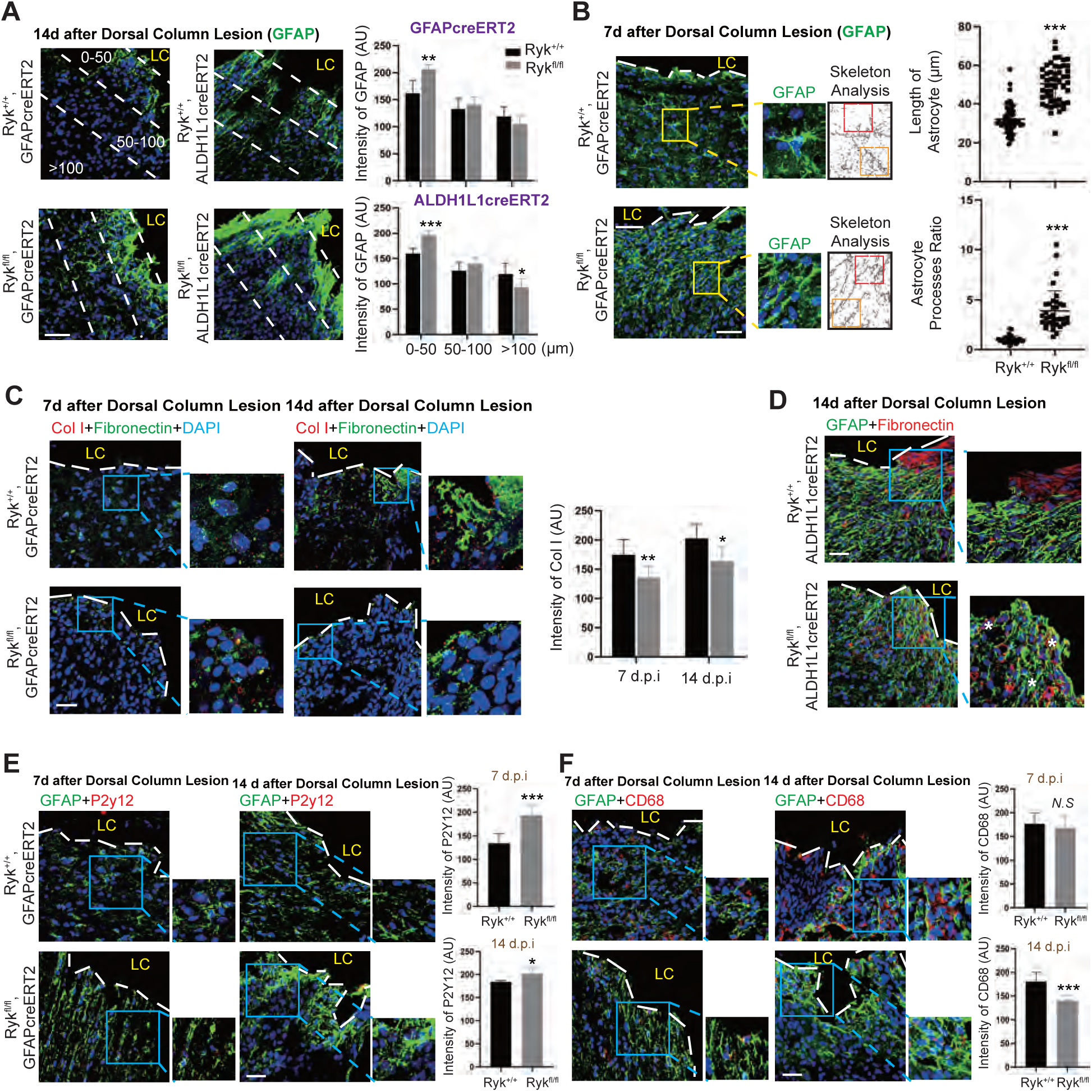
Accelerated astrocyte border formation, enhanced polarization and elongation of astrocyte processes, reduced fibrosis and accelerated inflammatory responses in astrocyte-specific Ryk conditional knockout. **(A)** Astrocyte border strained with GFAP antibody 14 days after SCI in control or *Ryk cKO.* Scale bar =40 µm. Bar graphs show GFAP staining at different distances from the lesion border. N=5 for each group. **(B)** Skeleton analysis for astrocytes around the lesion border 7 days after SCI. Astrocyte processes ratio: Length of processes toward the lesion site/ Length of processes on the other side. Scale bar =40 µm. N=5 for each group. **(C)** Staining of spinal cord sections with antibodies against Col1 or Fibronectin 7 days or 14 days after SCI. Scale bar =40 µm. Bar graphs showed quantitative analysis of Col I expression. **(D)** Staining of spinal cord sections with antibodies against GFAP and Fibronectin 14 days after SCI. The GFAP^+^ elongated astrocyte formed many ovoid-like structures surrounding fibroblasts (* labeling). Scale bar =40 µm. **(E)** Staining of spinal cord sections with antibodies against GFAP and P2Y12 7 days or 14 days after C5 dorsal column lesion in control and *Ryk cKO.* Scale bar =40 µm. N=3 for each group. **(F)** Staining of spinal cord sections with antibodies against GFAP and CD68 7 days or 14 days after C5 dorsal column lesion in control and *Ryk cKO* . Scale bar =40 µm. N=3 for each group. Data are expressed as mean ± SD. \**P* < 0.05, \*\**P* < 0.01 and \*\*\**P* < 0.001 vs. the indicated groups.

### Increased carrolling of fibroblasts by astrocytes and accelerated resolution of inflammatory responses of microglia in astrocyte-specific *Ryk* conditional knockout

In addition to the change of astrocyte morphology at the lesion border in *Ryk cKO*, we also observed changes in fibroblasts. We co-stained for Col I and fibronectin and found that, indeed, astrocyte-specific *Ryk cKO* did reduce immunoreactivity of Col I 7 days and 14 days after injury, whereas the level of fibronectin was unchanged (Fig. 3C, S5A and S5B). 14 days after injury, astrocytes formed a distinct boundary between the lesion core and the parenchyma in both WT and *Ryk cKO* (Fig. 3C). In the WT lesion core, we observed aggregated fibroblasts with strong fibronectin expression. However, in *Ryk cKO*, we found that the fibronectin staining was greatly reduced in the lesion core. Fibronectin^+^ cells located in the parenchyma (marked with stars) were surrounded by the elongated astrocyte processes (Fig. 3D). These results are consistent with the reported observation of carrolling of fibroblasts by astrocytes (8).

P2y12^+^ microglia are considered as a marker of homeostatic microglia. The immunostaining data showed that P2y12 expression in *Ryk GFAPcre ERT2 mice* was higher compared to WT mice both at 7 and 14 days after injury (Fig. 3E). And in *Ryk Aldh1l1creERT2* mice, P2y12 was found significantly increased at 7 days after injury while there was no significant difference 14 days after injury (Fig. S6A). These results suggest that there was an enhanced or accelerated microglia responses in astrocyte specific *Ryk cKO*. CD68^+^ microglia represent phagocytic microglia. Data showed that CD68 expression didn’t show significant difference 7 days after injury while a significant decrease in *Ryk cKO* 14 days after SCI (Fig. 3F and S6B). These data suggest that inflammation progressed faster and got resolved earlier in astrocyte-specific *Ryk* cKO.

### Single-cell transcriptomics analyses revealed extensive changes of cell-cell signaling in astrocyte-specific *Ryk* knockout

To explore how astrocyte-specific *Ryk cKO* promoted astrocyte polarization and astrocyte border formation and accelerated inflammatory responses, we performed single cell RNA sequencing (SCR seq) to identify potential changes of gene expression and cell-cell signaling among all cell types. We dissociated cells from the spinal cord tissue around the lesion core. Two biological repetitions of single cell suspensions were generated from the mice which uninjured, 1d, 7d, and 14d after SCI with or without *Ryk cKO* by *hGFAP-CreERT2.* After data quality control, a total of 18,203 high quality cells remained. Batch effects were corrected by Harmony. The cells were then subjected to the standard Seurat pipeline of normalization, feature selection, dimensionality reduction first by principal component analysis (PCA) and then uniform manifold approximation and projection (UMAP). Clustering was computed via Leiden algorithm in Seurat. 17 distinct cell clusters were resolved (Fig. 4A). According to the highest differentially expressed genes (DEGs) and canonical markers used in prior studies, we annotated these 17 clusters as 13 major cell types with distinct expression profiles (Fig. 4A, S7): astrocytes (ASC, containing 3 clusters), microglia (MG, containing 2 clusters), oligodendrocytes (containing 2 clusters), endothelial cells, pericytes, fibroblasts, neuronal cells, ependymal cells, blood cells and immune cells. We first looked into the astrocyte subtypes. The expression of the reactive astrocyte marker gene GFAP was the highest in Cluster 5 astrocytes (Fig. S8). Other markers Aqp4 (Aquaporin 4) and Gja1 (Connexin43) were highly expressed in Cluster 5 and Cluster 7 astrocytes but not in Cluster 2 astrocytes. And S100B, a marker for more mature grey matter astrocytes, was highly expressed in Cluster 2 and Cluster 5 astrocytes (13). Known border-forming astrocyte markers, Cdh2, Csgalnact1 and Chst11 were found expressed more in Cluster 7 (14). Therefore, Cluster 7 astrocytes are likely border-forming astrocytes. Cluster 2 may be another more differentiated subtype derived from reactive astrocytes (15).

**Fig. 4.**
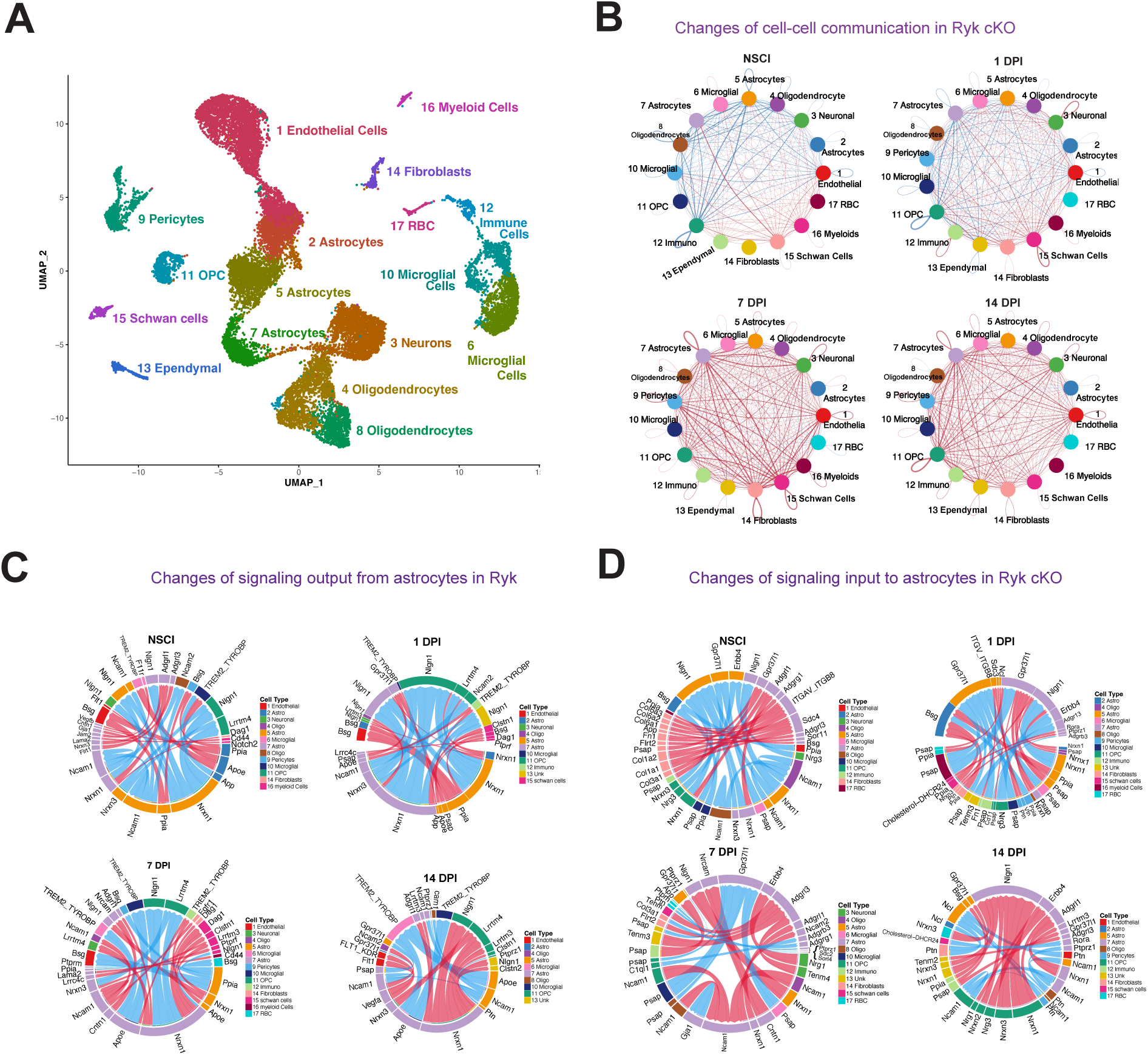
Predicted change of cell-cell signaling in astrocyte-specific *Ryk cKO* at the injury site after spinal cord injury. **(A)** UMAP plot of major cell types (18,203 cells). **(B)** CellChat chord diagram showing changes of strength of cell-cell signaling in *Ryk cKO*. Red lines represented increase in signaling strength while blue lines represented decrease. **(C)** Changes of signaling output from astrocytes in *Ryk cKO*. **(D)** Changes of signaling input to astrocytes in *Ryk cKO*.

To identify the potential changes of cell-cell signaling, we quantified and visualized the global communication atlas among all cell types using a computational tool, CellChat (16). CellChat predicted that spinal cord injury changed the expression of the components of a large number of signaling pathways over the course of 1 day, 7 days and 14 days after injury in the wildtype control, especially in fibroblasts, ependymal cells, immune cells, myeloid cells and microglia (Fig. S9A). These changes are greatly increased over the course of 1 day, 7 days and 14 days after injury in astrocyte-specific *Ryk cKO* (Fig. S9B). When we directly compared astrocyte specific Ryk conditional knockout with control, we found that the inferred strengths of interactions among all cell types are in general broadly increased in astrocyte specific *Ryk cKO*, especially after 7 days and 14 days after injury, particularly in Cluster 7 astrocytes, Cluster 6 microglia and fibroblasts (Fig. 4B). The greatest increase of signaling strength appears to occur on day 7 post-injury as indicted by the thickness of the red lines, which indicates strength (Fig. 4B). The lists of the pathways which are predicted to undergo increase or decrease due to astrocyte-specific Ryk cKO also suggest that astrocyte specific *Ryk cKO* profoundly changed many signaling pathways (Fig. S10). The greatest increase of signaling pathways in astrocyte specific *Ryk cKO* was observed 7 days post-injury (Fig. S10C). And 14 days post-injury, astrocyte-specific *Ryk cKO* led to a decrease of CPSG4 and MAG, which are well-known to inhibit axonal regeneration, suggesting that astrocyte-specific *Ryk cKO* may lead to a more permissive environment for axon regeneration (Fig. S10D). Another important signaling pathway is VGEF, which promotes angiogenesis. We found that VGEF signaling was initially down regulated on 1 day post-injury in astrocyte-specific *Ryk cKO* but became increasingly up regulated on 7 days and 14 days post-injury between astrocytes and endothelial cells and between astrocytes and pericytes (Fig. S11).

To understand how *Ryk cKO* may lead to the dramatic morphological changes of astrocytes, we looked into the cell adhesion pathways. We noticed that the ligand-receptor pair Cntn1-Nrcam was unchanged in astrocyte-specific Ryk knockout in uninjured animals but was ranked on of the top increased 7 days after injury, suggesting that NrCAM, a transcriptional target of β-catenin/LEF-1 pathway, may be subject to the regulation of Ryk only in astrocytes injury settings (17) (Fig. 4C). The Psap-GPR37/GPR37L1 neuroprotective and glioprotective signaling was predicted to be among the top increased pathways 14 days post-injury in astrocytes, again suggesting that astrocyte-specific *Ryk* knockout promotes neuroprotection and regeneration (18) (Fig. 4D).

### Increased pro-regenerative signaling and accelerated astrocyte differentiation in astrocyte-specific *Ryk* knockout

The extensive changes of cell-cell signaling pathways, which are highly relevant to neuroprotection and regeneration prompted us to analyze canonical Wnt signaling, which is essential for activating genes for regeneration and repair after spinal cord injury and can also affect many other signaling pathways (19–22). Ryk is a co-receptor of Frizzled for Wnt and may be a regulator of Wnt signaling in astrocytes after spinal cord injury. CellChat analyses predicted that canonical Wnt signaling pathway was significantly increased from astrocytes to fibroblasts and endothelia cells as well as from fibroblasts to endothelial cells and fibroblasts to fibroblasts themselves in astrocyte-specific *Ryk cKO* at 7 days after injury. 14 days after injury, canonical Wnt signaling was significantly increased among all these cells as well as from astrocytes to OPC (Fig. 5A). To validate these findings, we stained for phosphorylated β-catenin and found that it is strong, around 2 fold, increased in the nuclei of astrocytes in *Ryk cKO* 7 days after injury (Fig. 5B). Interestingly, phosphorylated β-catenin levels in control and *Ryk cKO* are similar 14 days after injury, suggesting that the loss of *Ryk* in astrocytes lead to a transient increase of canonical Wnt signaling (Fig. 5C). Consistent with this, we only observed an increase of BrdU incorporation in astrocytes in *Ryk cKO* 7 days but not 14 days after injury (Figures S4C and S4D). FGF signaling, which regulates glial cell morphology and also implicated in repair after injury, was significantly increased in astrocytes and between astrocytes and many other cell types after injury in astrocyte-specific *Ryk cKO* (Fig. 5D) (23) (24) (25). Therefore, the two key signaling essential for injury response and repair are dynamically altered among multiple cell types.

**Fig. 5.**
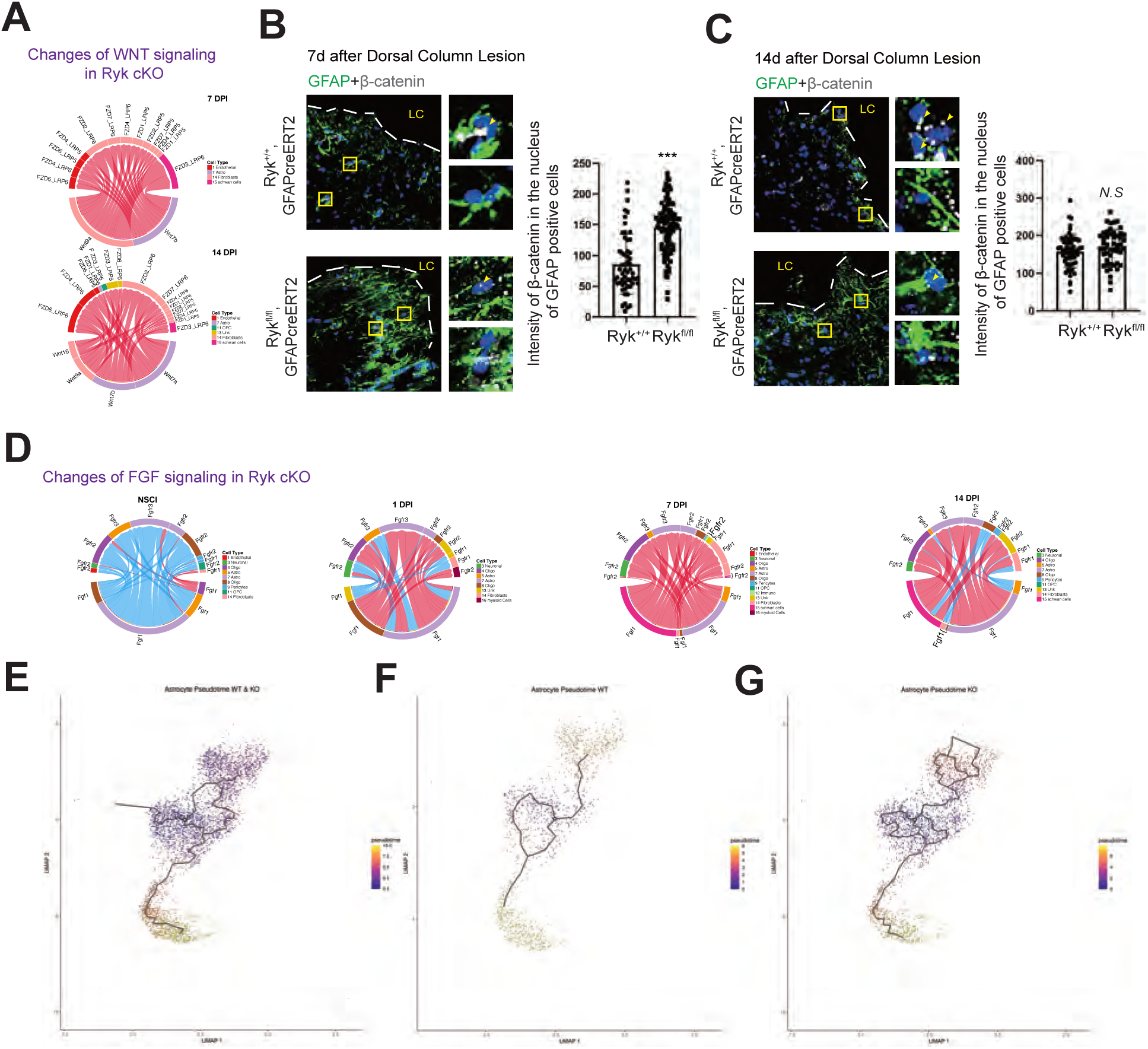
Changes of Wnt and FGF signaling and astrocyte trajectory astrocyte-specific *Ryk cKO*. (**A**) CellChat analyses predict a strong increase of canonical Wnt signaling involving astrocytes astrocyte-specific *Ryk cKO*. (**B**) Immunostaining of spinal cord sections with GFAP and phosphorylated β-catenin 7 days after injury. Scale bar =40 µm. Dot graphs showed quantitative analysis of phosphorylated β-catenin expression in astrocyte nuclei. Data are expressed as mean ± SD. \*\*\**P* < 0.001 vs. the indicated groups. **(C)** Immunostaining of spinal cord sections with GFAP and phosphorylated β-catenin 14 days after injury. Scale bar =40 µm. Dot graphs showed quantitative analysis of phosphorylated β-catenin expression in astrocyte nuclei. **(D**) CellChat analyses showing the changes of FGF signaling in astrocyte-specific *Ryk cKO..* **(E)** Pseudotime analyses of astrocyte trajectory in control and *Ryk cKO*. **(F)** Pseudotime analyses of astrocyte trajectory in control. **(G)** Pseudotime analyses of astrocyte trajectory in *Ryk cKO*.

The broad changes of cell signaling prompted us to ask more systematically whether the progression of astrocyte differentiation is altered by cell trajectory analysis (26, 27). As GFAP is a marker for reactive astrocytes and S100B (high in type 2) and Cdh2 (border-forming astrocyte marker high in type 7) are markers for more differentiated astrocytes, we selected type 5 as the start point for trajectory analyses (Fig. 5E and Fig. S8B). *Ryk cKO* significantly increased the number of type 2 and type 7 astrocytes, suggesting a faster maturation of astrocytes in *Ryk cKO* (Fig. 5F and 5G).

### Single-cell transcriptomics indicated accelerated inflammatory responses and pro-regenerative shift of microglia in astrocyte-specific conditional *Ryk* knockout

Microglia also play an important role in astrocyte border and fibrotic scar formation. Studies showed that appropriate microglia activation could remove harmful substances during injury and maintain homeostasis (28, 29). We found that expression of P2Y12, a marker for parenchymal M2 pro-regenerative microglia, in microglia was increased from 7 days post injury or earlier but expression of CD68, a marker for pro-inflammatory phagocytic microglia, was reduced 14 days post-injury in astrocyte-specific conditional *Ryk* knockout (Fig. 3E-3F and Fig. S6) (30–33). The fact that microglia responded to astrocyte-specific conditional *Ryk* knockout at an earlier time point than the formation of glial border suggests that the accelerated microglia response may not be caused by reduced tissue damage due to faster closure of the glial border. Therefore, we analyzed the responses of microglia in our single cell sequencing data. CellChat analysis data showed that, indeed, the ApoE-TREM2_TYROBP pathway was a top decreased pathway in microglia by Day 14, although it was initially increased on Day 1 (Fig. 6A, 6B). Similar to astrocytes, we performed pseudotime analyses in microglia. We found that Cluster 10 microglia have lower expression of P2Y12 but higher CD68, whereas Cluster 6 has higher P2Y12 and lower CD68 (Fig. S8B). Therefore, Cluster 6 may represent the pro-regenerative M2 type of microglia and Cluster 10 may be the pro-inflammatory M1 type. We found that astrocyte-specific conditional *Ryk* knockout shifted the trajectory from Cluster 10 to Cluster 6 and increased the frequency of the more differentiated states (as indicated by higher proportion in the pseudotime bins in stages 9 and 10) (Fig. 6C-6F).

**Fig. 6.**
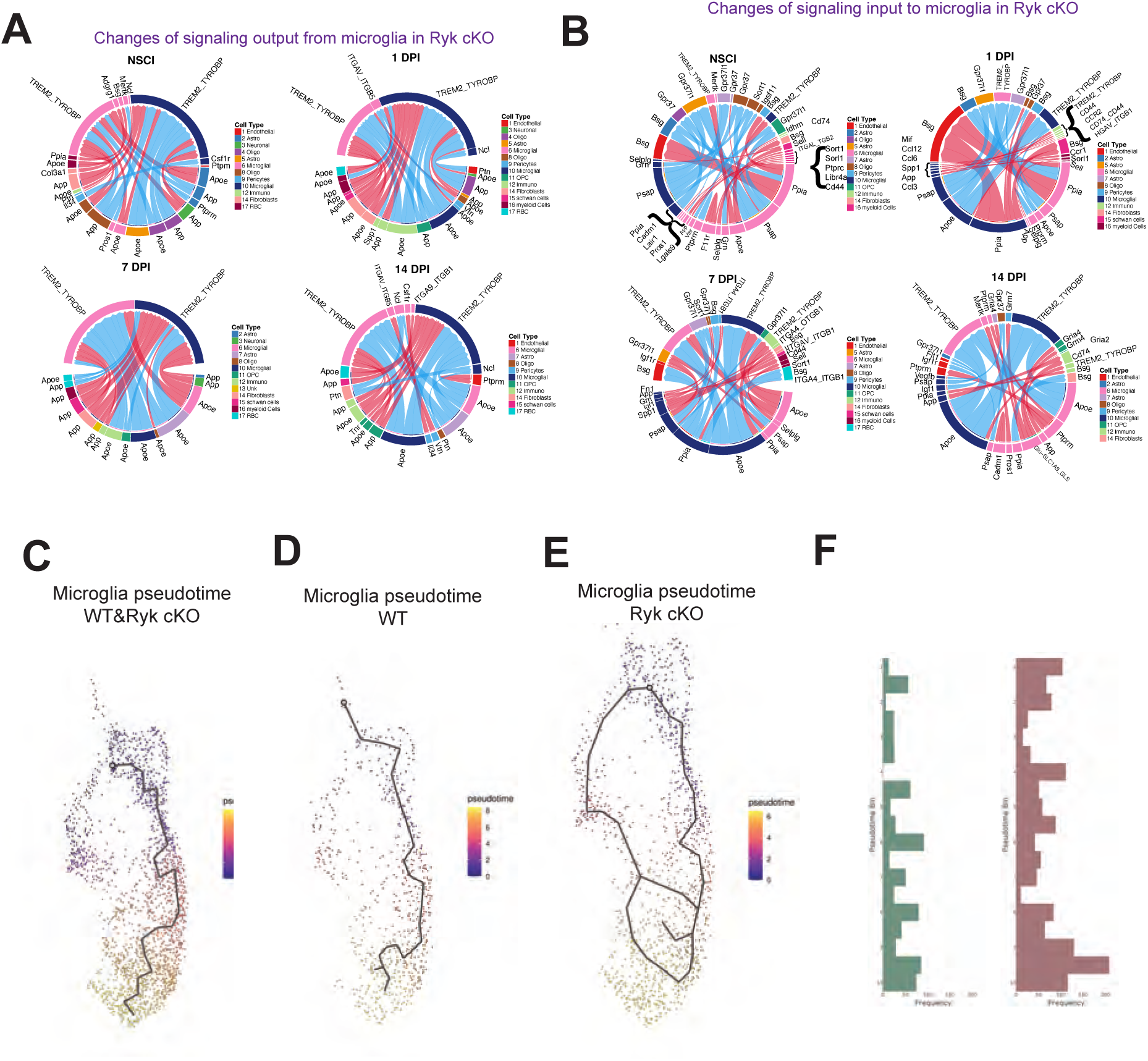
Cell signaling changes in microglia in astrocyte-specific Ryk cKO after SCI. **(A)** Changes of signaling output from microglia in *Ryk cKO*. **(B)** Changes of signaling input to microglia in *Ryk cKO*. **(C)** Pseudotime analyses of microglia trajectory in control and *Ryk cKO*. **(D)** Pseudotime analyses of microglia trajectory in control. **(E)** Pseudotime analyses of microglia trajectory in *Ryk cKO*. (**F**) Histogram showing the number of cells in bins that represent pseudotime trajectory (from 1 to 10).

### NrCAM is required for the elongation of the processes of reactive astrocytes and formation of glial border

To test our hypotheses based on our spinal cord injury results (Fig. 1-3) and single cell transcriptomics analyses (Fig. 4-6), we first examined NrCAM expression in astrocytes using immunostaining. We found that there was almost no NrCAM immunoreactivity in astrocytes 7 days after injury in WT mice. However, NrCAM staining was significantly increased in astrocytes in *Ryk cKO* (Fig. 7A and S12A). 14 days after SCI, the NrCAM expression level in astrocyte started to increase in the area immediately abutting the lesion core in WT mice. And the NrCAM expression in Ryk cKO mice is still higher than WT (Fig. 7B and S12B). These data suggested that Ryk cKO might accelerate astrocyte border formation by up-regulating NrCAM signaling in astrocytes.

**Fig. 7.**
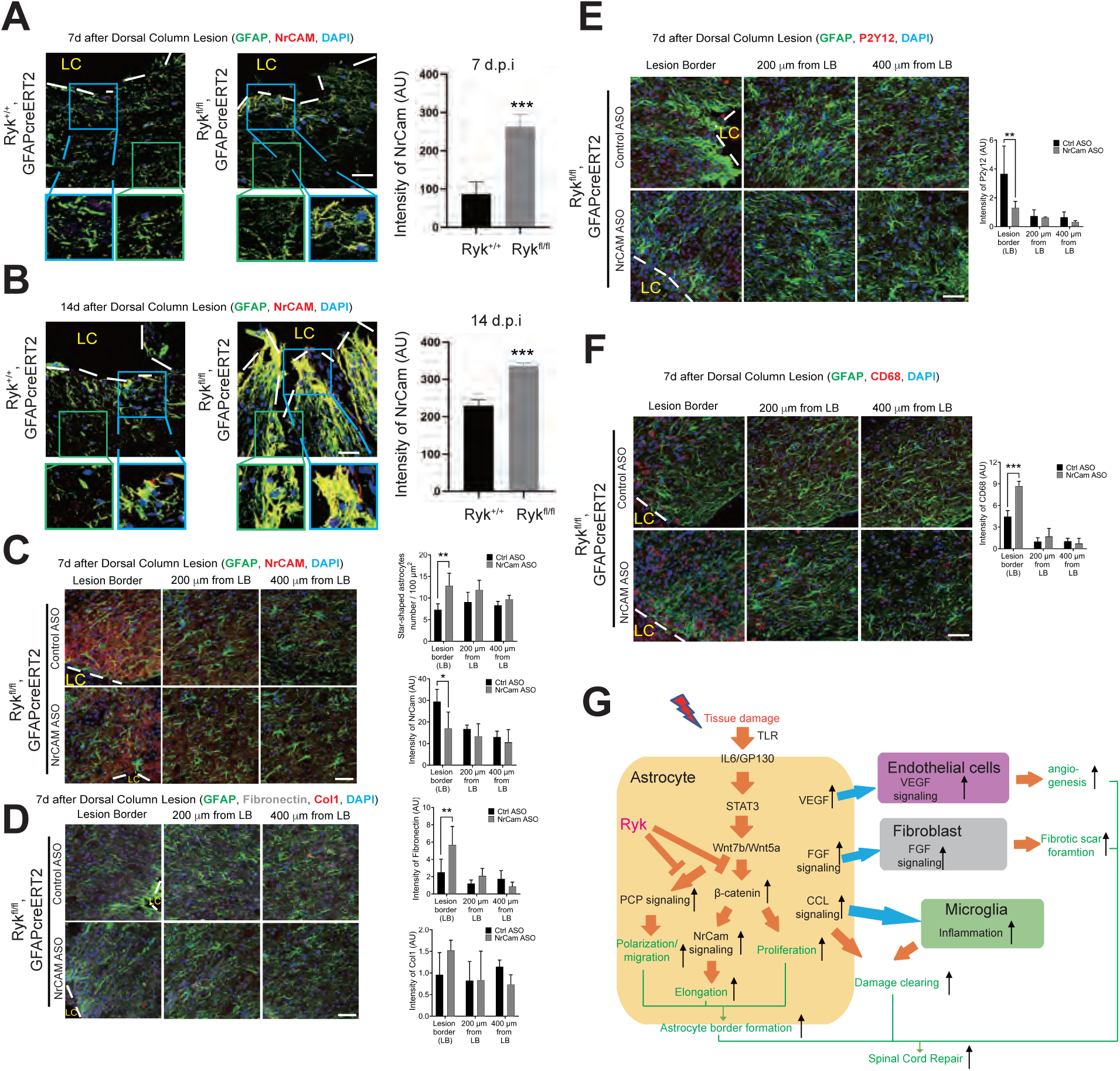
NrCAM is required for the elongation of the processes of border-forming reactive astrocytes. (**A**) Immunostaining of spinal cord sections with antibodies against NrCAM and GFAP in control or *Ryk cKO* 7 days after injury. Scale bar =40 µm. N=3 for each group. Blue box labelled the areas immediately abutting the lesion core while green box was about 100 µm away from the lesion core. Bar graphs showed quantitative analysis of the intensity of NrCAM. **(B)** Immunostaining of spinal cord sections with antibodies against NrCAM and GFAP in control or *Ryk cKO* 14 days after injury. Scale bar =40 µm. N=3 for each group. Blue box labelled the areas immediately abutting the lesion core while green box was about 100 µm away from the lesion core. Bar graphs showed quantitative analysis of the intensity of NrCam. **(C)** Immunostaining of spinal cord sections with antibodies against NrCAM and GFAP to analyze the number of star-shaped astrocytes and level of NrCAM in *Ryk cKO* 7 days after C5 dorsal column lesion and ASO injection. N=3 for each group. **(D)** Immunostaining of spinal cord sections with antibodies against fibronectin, Col1 and GFAP to analyze the level of Fibronectin and Col1in *Ryk cKO* 7 days after C5 dorsal column lesions and ASO injection. Scale bar =40 µm. N=3 for each group. **(E)** Immunostaining of spinal cord sections with antibodies against P2Y12 and GFAP to analyze the level of P2Y12 in *Ryk cKO* 7 days after C5 dorsal column lesion and ASO injection. Scale bar =40 µm. N=3 for each group. **(F)** Immunostaining of spinal cord sections with antibodies against CD68 and GFAP to analyze the level of CD68 in *Ryk cKO* 7 days after C5 dorsal column lesion and ASO injection. Scale bar =40 µm. N=3 for each group. (**G**) Summary of Ryk function in astrocyte border formation, astrocyte differentiation and cell-cell signaling to coordinate multi-cellular interactions in wound healing. Data are expressed as mean ± SD. \**P* < 0.05, \*\**P* < 0.01 and \*\*\**P* < 0.001 vs. the indicated groups.

To further test this hypothesis, we designed antisense oligos (ASO) against NrCAM to reduce its expression. After detecting the efficiency of ASOs by RT PCR, we injected two NrCAM ASOs (separately) into the parenchyma of the spinal cord near the injury site immediately after SCI and obtained similar results (Fig. S13). The immunostaining results verified that NrCAM ASO injection significantly reduced the expression of NrCAM in astrocytes near the lesion core in Ryk cKO 7 days after injury (Fig. 7C). NrCAM ASO reversed Ryk cKO-induced astrocyte elongation and polarization at 7 days after injury and star-shaped astrocytes, which are rare at the lesion border, were significantly increased (Fig. 7C). We then further tested fibrosis and microgliosis and found that the *Ryk cKO* induced changes (Fig. 3C-3F) were both reversed in animals treated with NrCAM ASO (Fig. 7D-7F). The above data suggested that *Ryk cKO* accelerated astrocyte border formation by up-regulating cell adhesion pathways in astrocytes, which may be critical for the changes of the morphology of the astrocytes. Among them, NrCAM is a major player.

## DISCUSSION

The complex cellular interactions for wound healing after central nervous system injury is poorly known. Reactive astrocytes play pivotal roles in injury responses in glial border formation. We uncovered a major signaling pathway, downstream of STAT3, which regulates multiple aspects of astrocyte responses (34–36) (Fig. 7G). We propose that the canonical Wnt signaling pathway, once activated by STAT3, activates astrocyte proliferation, differentiation (increases of thickness and elongation of astrocyte processes) and inflammatory responses (for tissue debris clearing).

Ryk, a Wnt receptor, negatively regulates canonical Wnt pathways in astrocytes in this context. In astrocyte-specific Ryk conditional knockout, we found that the expression of a direct transcriptional target of β-catenin, NrCAM, was greatly increased and NrCAM is essential for the elongation of astrocyte processes and potentially migration of astrocytes (17). Therefore, we identified a complete transcriptional cascade from STAT3 to NrCAM, via canonical Wnt β-catenin signaling pathway, to regulate reactive astrocyte morphology for glial border formation after spinal cord injury.

How astrocytes coordinate signals to other cell types for wound healing is not well understood. Our study suggests that Ryk may coordinate many signals emanating from astrocytes to coordinate responses of many cell types after spinal cord injury (Fig. 7G). VEGF, which promotes angiogenesis, is also a direct transcriptional target of canonical Wnt signaling pathway and its expression was found increased in astrocyte-specific Ryk conditional knockout. The large increase of FGF signaling from astrocytes to fibroblasts in *Ryk cKO* may lead to faster fibroblast proliferation and wound healing, resulting in less inflammation. The accelerated inflammatory responses of both astrocytes and microglia may result in more efficient tissue debris clearing and tissue repair. Because wound healing and tissue repair involve the interactions among many cell types, it is important to coordinate the proliferation and differentiation of all cell types involved. We propose that the normal function of Ryk may be to control the rate of cellular responses to allow time for such coordination. However, it is possible that injury may over activates this function of Ryk. Therefore, inhibiting Ryk function may benefit wound healing.

Our study also revealed a potential coordinated polarization of astrocytes and axons by the same Wnt/Ryk signaling mechanism. Ryk is a Wnt receptor and regulates a non-canonical Wnt pathway, the planar cell polarity (PCP) pathway, in growth cone guidance and synapse maintenance in neurons (11, 37, 38). Whether the PCP pathway is also active in astrocytes to regulate the polarization of astrocytes towards the lesion site awaits future studies. If this is the case, the induced Wnts at the lesion site may prevent astrocytes from polarizing towards and/or migrating to the lesion border in similar ways that the axons are repelled away from the injury site (2). Therefore, inhibiting Ryk function in astrocytes can allow astrocytes to be better polarized towards and migrate to the lesion border to form scar border more efficiently to accelerate the glial border formation.

By conditionally knocking out *Ryk* in astrocytes, we were able to accelerate the formation of glial border and reduce the lesion volume. Our analyses showed that this accelerated astrocyte border formation is beneficial for preserving and recovery sensory-motor functions probably by better protection of the spinal cord tissues due to the faster healing and reduced secondary injury. Our previous work showed that intrathecal infusion of a function blocking Ryk antibody in injured spinal cord resulted in greater functional recovery than conditional knocking out Ryk from the corticospinal tract (CST) neurons in a skilled motor task (5). Given our new findings here, we propose that the additional functional recovery may be caused by the faster astrocyte border formation in addition to promoting the growth of CST axons. Therefore, targeting Ryk may have at least two benefits, promoting axon growth and accelerate astroglial border formation to protect spared spinal cord tissue, including neurons. We observed that in injured human spinal cord tissues, Ryk was also induced in both astrocytes and the injured axons, suggesting that Ryk is a promising therapeutic target to treat human spinal cord injury (Fig. 1C-1E).

## MATERIALS AND METHODS

### Animals

All animal work in this research was approved by the University of California, San Diego (UCSD) Institutional Animal Care and Use Committee. Animals were housed on a 12 h light/dark cycle and behavioral analyses were done at consistent morning hours during the light cycle. GFAPcre^ERT2^ (Strain #:012849) and Aldh1L1 cre^ERT2^ (Strain #:029655) mice were purchased from the Jackson Laboratory. Ryk^fl/fl^ (cKO) was generated in the Zou laboratory. Mice aged 8-10 weeks were be used in the following experiments.

### Surgical procedures

#### C5 dorsal column lesion

Mice were deeply anaesthetized with ketamine, spinal level C5 was exposed by laminectomy and the dorsal columns were lesioned at a depth of 1 mm with Vannas spring scissors (Fine Science Tools, Foster City, CA). The dorsal musculature was sutured with 5-0 silk sutures and the skin was closed with wound clips.

#### NrCam antisense oligo design and injection

NrCam ASOs are designed and produced by IDT. We designed 3 ASOs against NrCam: NrCam- 1 (C*C*A*C*G*C*T*G*A*C*G*C*G*A*A*C*A*T*T*T) NrCam-2 (C*T*G*T*C*G*T*G*C*G*T*G*T*T*T*C*C*G*A*A) and NrCam-3 (G*A*C*G*G*C*T*C*C*T*A*A*T*G*C*G*T*T*T*T). Negative Control ASO: G*C*G*A*C*T*A*T*A*C*G*C*G*C*A*A*T*A*T*G. After efficiency testing by RT-PCR, we used NrCam-2 and -3 for subsequent animal experiments.

ASO injection was performed as described previously. Briefly, immediately after lesion, two injections of ASO were made into the spinal cord along the dorsal midline at 0.5 mm rostral and 0.5 mm caudal to the lesion site. For each injection site, two injections at depths of 0.5 and 1.0 mm were made (0.5 µl at each depth) using a pulled glass capillaries attached to a pneumatic Pico Pump (World Precision Instruments).

#### Corticospinal Tract Tracing

Corticospinal Tract Tracing was performed as we described previously. Briefly, mice were anesthetized and stabilized in a stereotaxic frame. Using a digital stereotactic injector (Item: 51709, stoelting Co. USA), 0.5 µl of biotin dextran amine (BDA; MW 10,000; 10% in PBS; Molecular Probes) was injected into one of the 10 total sites (5 sites/side). Mediolateral (ML) coordination: 1.5 mm lateral to the bregma; anteroposterior (AP) coordination from the bregma: 1.0, 0.5, 0, 0.75 and 1.5 mm; dorsoventral (DV) coordination: 0.5 mm from the cortical surface. After each injection was completed, the injector tip was left in place for an additional 5 min to ensure that the BDA solution adequately penetrated the tissue. Two weeks later, mice were anesthetized and perfused with 4% paraformaldehyde for detecting CST distribution in the spinal cord.

### Behavioral Tests

#### Grip strength tests

Grip strength was measured with a computerized grip strength meter (Bio-GS3, Bioseb, USA). To measure grip strength in the forepaws of the mice, the experimenter held the mice gently by the base of the tail, allowing the animal to grasp the grid with the forepaws. As soon as the mice grasped the transducer metal grid with their forepaws, the experimenter pulled the animals backwards by the tail until grip was lost. The peak force of each measurement was automatically recorded in grams (g) by the device. Forelimb grip strength in each mouse was measured twice. Rotarod performance To measure the balance and ability to coordinate stepping, mice were placed on a single lane rotarod (Med Associates) for two trials per session. The rotarod was set for constant acceleration from 3.0 to 30 rpm over 300 s and mice were scored on seconds to fall. The final result is the average of the two trials.

### Immunohistochemistry

Mice were sacrificed and were perfused with saline solution, followed by 4% paraformaldehyde in 0.1 M phosphate-buffered saline. The spinal cords were then dissected out, fixed in 4% paraformaldehyde overnight and equilibrated in 30% sucrose at 4°C. Sagittal sections (30 µm) were cut on a freezing microtome (Thermo, CRYOSTAR NX50). After rinsing with 0.1 % Triton X-100 (vol/vol; Sangon, T0694) in 0.1 M PBS and blocking with blocking buffer (1% bovine serum albumin and 5% donkey serum in TBS solution with 0.1% Triton X-100) for 1 h at room temperature, sections were incubated with the following primary antibodies overnight at 4°C: rabbit anti-Ryk (generated in the Zou laboratory), chicken anti-GFAP (Abcam, ab4674), rabbit anti-GFAP (Abcam, ab7260), mouse anti-GFAP (Sigma, G3893-100UL), mouse anti-fibronectin (Millipore, CP70), rabbit anti-Col 1A1 (Novus Biologicals, NBP1-30054), rabbit anti-NrCam (Abcam, ab24344), rabbit anti-CD68 (Abcam, ab125212),rabbit anti-p2y12 (Invitrogen, PA5-77671) and mouse anti-NeuN (Abcam, ab104224). Afterward, sections were incubated with Alexa Fluor 647-, 594- or Alexa Fluor 488-conjugated secondary fluorescent antibody (1:400; Jackson ImmunoResearch) for 1 h at room temperature, counterstained with 4′ diamidino-2-phenylindole (DAPI), and mounted in mounting media. Images were captured using a Zeiss 880 Airyscan microscope.

### Immunostaining for human spinal cord samples

The slides were microwaved slides in Antigen Retrieval Buffer (2 mins at 90% power, 2 mins at 70% power, and 6 mins at 50% power). The slides were then removed from microwave and incubated at room temperature in Antigen Retrieval Buffer for 20 mins and rinsed in 1X TBST at room temperature. Endogenous peroxidase was blocked with 3% H2O2 in 50% methanol for 30 mins. After rinsing with 1X TBST and blocking with blocking buffer for 0.5 h at room temperature, sections were incubated with the following primary antibodies overnight at room temperature: mouse anti-Ryk (5), chicken anti-GFAP (Abcam, ab134436), rabbit anti-Neurofilament (Sigma, N4142) and rabbit anti-SOX9 (Abcam, ab185230). After rinsing with 1X TBST, sections were incubated with Alexa Fluor 647-, 594- or Alexa Fluor 488-conjugated secondary fluorescent antibody (1:400; Jackson ImmunoResearch) for 1 h at room temperature. After rinsing, slides incubated in 95% ethanol for 1 min at room temperature, in 100% ethanol for 1 min*3 times at room temperature and in xylene for 1 minute*3 times at room temperature. Slides were then incubated with 4′,6-diamidino-2-phenylindole (DAPI) for 15 min, and mounted in mounting media. Images were captured using a Zeiss 880 Airyscan microscope.

### Single cell RNA seq

Mice were anesthetized and were performed a trans-cardiac perfusion with fresh ice-cold carbogen-bubbled cutting solution (212 mM sucrose, 1.25 mM NaH2PO4, 26 mM NaHCO3, 10 mM Glucose, 3 mM KCl, 7 mM MgSO4, 0.5 mM CaCl2). After decapitation, the spinal cords were immediately extracted and placed in fresh ice-cold cutting solution bubbled with a carbogen gas (95% O2 and 5% CO2). And were quickly transferred to the vibratome and sectioned into 300 μm slice. The slices were incubated in ACSF (125 mM NaCl, 1.25 mM NaH2PO4, 26 mM NaHCO3, 10 mM Glucose, 2.5 mM KCl, 1.3 mM MgSO4, 2 mM CaCl2) continuously aerated with carbogen gas.

Spinal cords were cut into small pieces < 1 mm in each dimension by a knife and collected to a 60 mm Petri dish leaving only enough ACSF to cover the tissues. The tissues were digested by pronase solution (1 mg/mL pronase, 45 μM ActD, 100 μg/mL DNase I in ice-cold carbogen-bubbled ACSF) at room temperature for 60 min with gentle agitation. After digestion, the tissues were exchanged into trituration buffer (1% FBS, 3 μm ActD, 100 μg/mL DNase I in ice-cold carbogen-bubbled ACSF) and triturated very gently through fire polished salinized Pasteur pipettes with the opening of 600 μm and 300 μm diameter.

The supernatant was collected and centrifuged at 300 rcf for 3 min at 4 ℃. The cell pellet was resuspended in wash buffer (0.1% BSA, 3 μm ActD, 100 μg/mL DNase I in ice-cold carbogen-bubbled ACSF) and filtered by 30 μm cell strainer. The dead cells were removed by MACS Dead Cell Removal Kit by manufactory’s instruction. The final single cell suspensions were rinsed in cold DPBS solution (0.1% BSA, 3 μm ActD, 100 μg/mL DNase I in ice-cold DPBS). The cell concentration and cell viability were determined by Cell Counter with AOPI staining. In our experiment, the cell viability of all samples was at least 80%. All steps were performed on ice or at 4 ℃ except the pronase digestion step. The wide-bore pipette tips were used to resuspend the cell pellet in all steps.

Fastqs were processed by 10X Cell Ranger and aligned to mm10 generated by Cell Ranger mkref. Each barcoded matrix from 10X cell ranger was loaded into R (V 4.1.2) and Seurat (V 4.3.0.1) and read with the Read10X function from Seurat. The percentage of mitochondria genes were used as quality control parameters for each cell. The thresholds used for a high-quality cell was percent mitochondrial genes < 25 and nFeature_RNA > 200. Seurat’s native log normalization was applied to the data and FindVariableFeatures function was used to find top 2000 variable genes used later for the ScaleData function. PCA was run to find 20 PCs which were then corrected using Harmony (V 0.1.1) package accounting for batch and sample effects. Using the corrected PCs from harmony, UMAP was run using Seurat’s RunUMAP and clusters were determined by the FindNeighbors and FindClusters function with a resolution parameter of 0.5 and using the Leiden algorithm revealing 17 clusters. To identify the cell-type of the clusters Seurat’s FindAllMarkers using default parameters was used (Wilcox ranked sum excluding genes with logFC < 0.25 and percent expression < 0.1). Further, a mixture of logFC and percent expression sorting by each cluster was used to identify marker genes while filtering for significant genes (p < 0.05). The top marker genes were compared to known marker genes for each cluster.

All 17 clusters were used for Cell Chat analysis (16). For each experimental group a Cell Chat object was created where default parameters were used for the following Cell Chat functions in this order: identifyOverExpressedGenes, identifyOverExpressedInteractions, computeCommunProb, computeCommunProbPathway, netAnalysis_computeCentrality, and aggregateNet. To compare differences between cell chat objects, mergeCellChat function was used and rankNet function was used to choose pathways of interest. To visualize pathways a modified netVisual_chord_gene function was used where Down-regulated edges were graphed as blue and upregulated edges were graphed as red in the chord diagram.

### Statistical analysis

All data were collected and analyzed in a blind manner. Data are presented as mean ± SD. One-way analysis of variance with least significant difference or Dunnett’s T3 post hoc test (where equal variances were not assumed) was applied for multiple comparisons, whereas Student’s t-test was used for comparisons between two groups. P < 0.05 was considered statistically significant.

## List of Supplementary Materials

Fig S1 to S13 for multiple supplementary figures

## Acknowledgments

We thank members of the Zou Lab for critical reading of the manuscript. Slide Scan and Airyscan confocal imaging were performed at UCSD School of Medicine Light Microscopy Facility (Grant P30 NS047101).

## Author contributions

Y.Z., Z.S. and B. F. designed the experiments; Z.S., B.F., W.L.,Y.L., S.V. and J.W. performed all experiments under the supervision of Y.Z; Z.S., B.F., W.L.,T.W. and Y.Z. analyzed and interpreted data; B.K.K. provided injured human spinal cord tissues. Z.S. and Y.Z. wrote the paper.

## Competing interests

Y.Z. is the founder of VersaPeutics and has equity, compensation and interim managerial role.

The terms of this arrangement have been reviewed and approved by the University of California,

San Diego in accordance with its conflict-of-interest policies. A disclosure of the design and sequences of antisense oligos against NrCAM has been submitted to UCSD Office of Innovation and Commercialization.

## Data and materials availability

The scRNA-seq raw and processed data generated in this study has been deposited in the NIH GEO database under accession code #GSE205029 (GEO Accession viewer (https://www.ncbi.nlm.nih.gov/geo)).

Otherwise, all data are available in the main text or the supplementary materials.

---

**Sup Fig 1.**
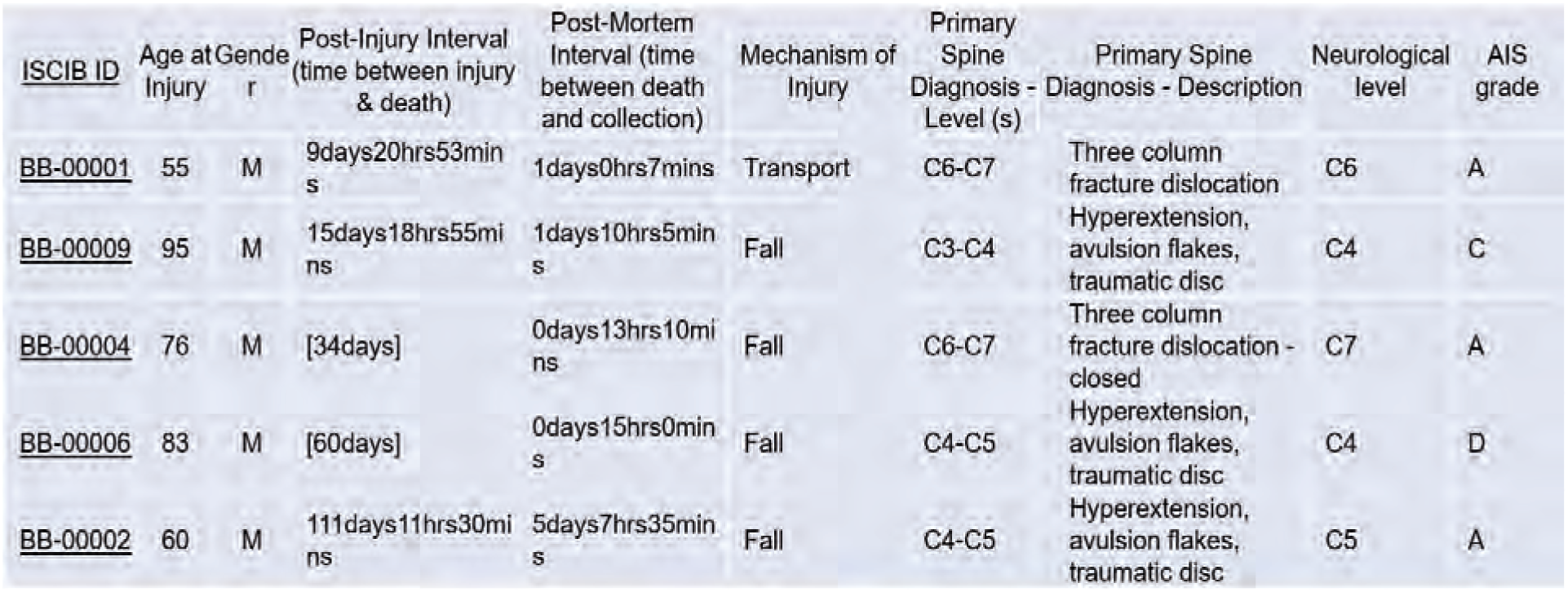
**Clinical information of spinal cord injury patients whose tissues were analyzed.**

**Sup Fig 2.**
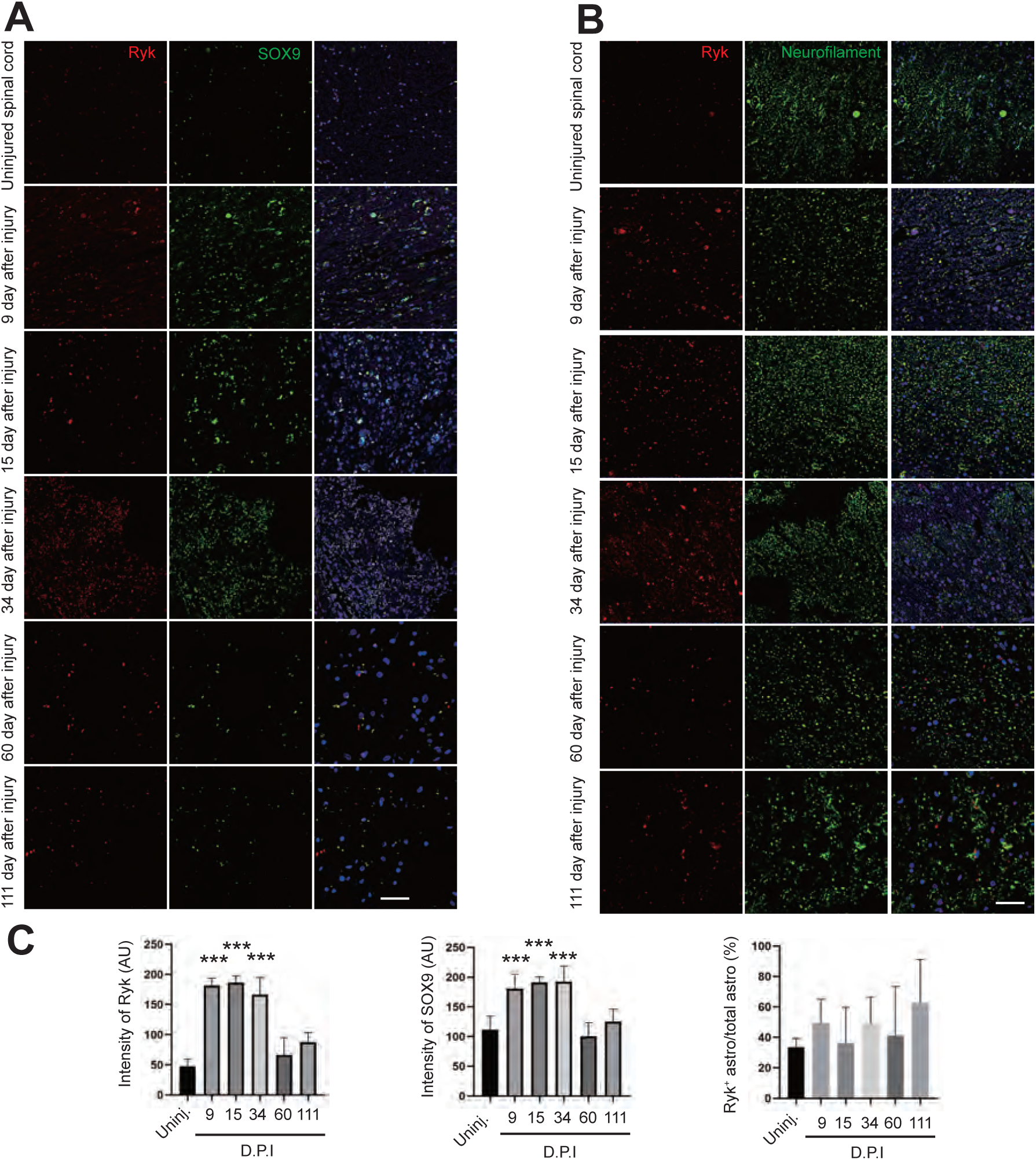
Induction of Ryk expression in human spinal cord injury away from the lesion epicenter. **(A)** Immunostaining of human spinal cord tissue (E4) at different time points after injury with antibodies against Ryk and SOX9. **(B)** Immunostaining of human spinal cord tissue (E1) at different time points after injury with antibodies against Ryk and Neurofilament H. **(B)** Bar graphs showing quantifications of Ryk expression level, SOX9 expression level and the percentage of Ryk^+^ astrocytes. Data are expressed as mean ± SD. \*\*\**P* < 0.001 vs. the indicated groups. Scale bar =40 µm.

**Sup Fig 3.**
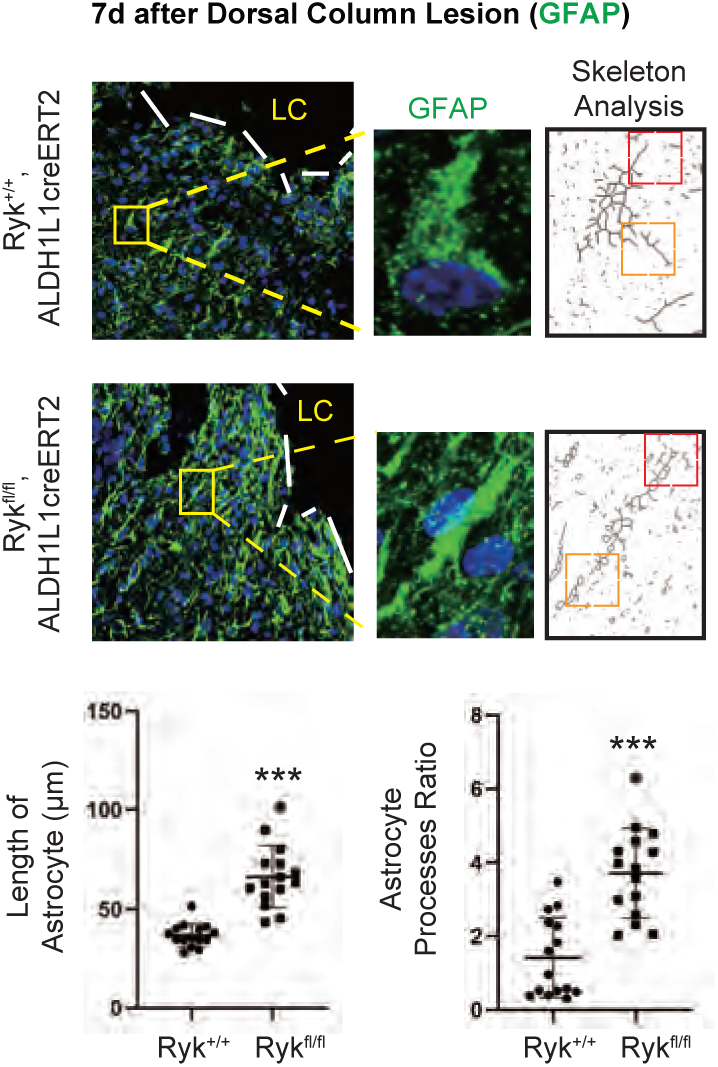
Increase of length and polarization of astrocyte processes in astrocyte-specific *Ryk cKO*. **(A)** Skeleton analysis for astrocytes around the lesion core 7 d after SCI. Scale bar =40 µm. N=5 for each group. \*\*\**P* < 0.001 vs. control groups.

**Sup Fig 4.**
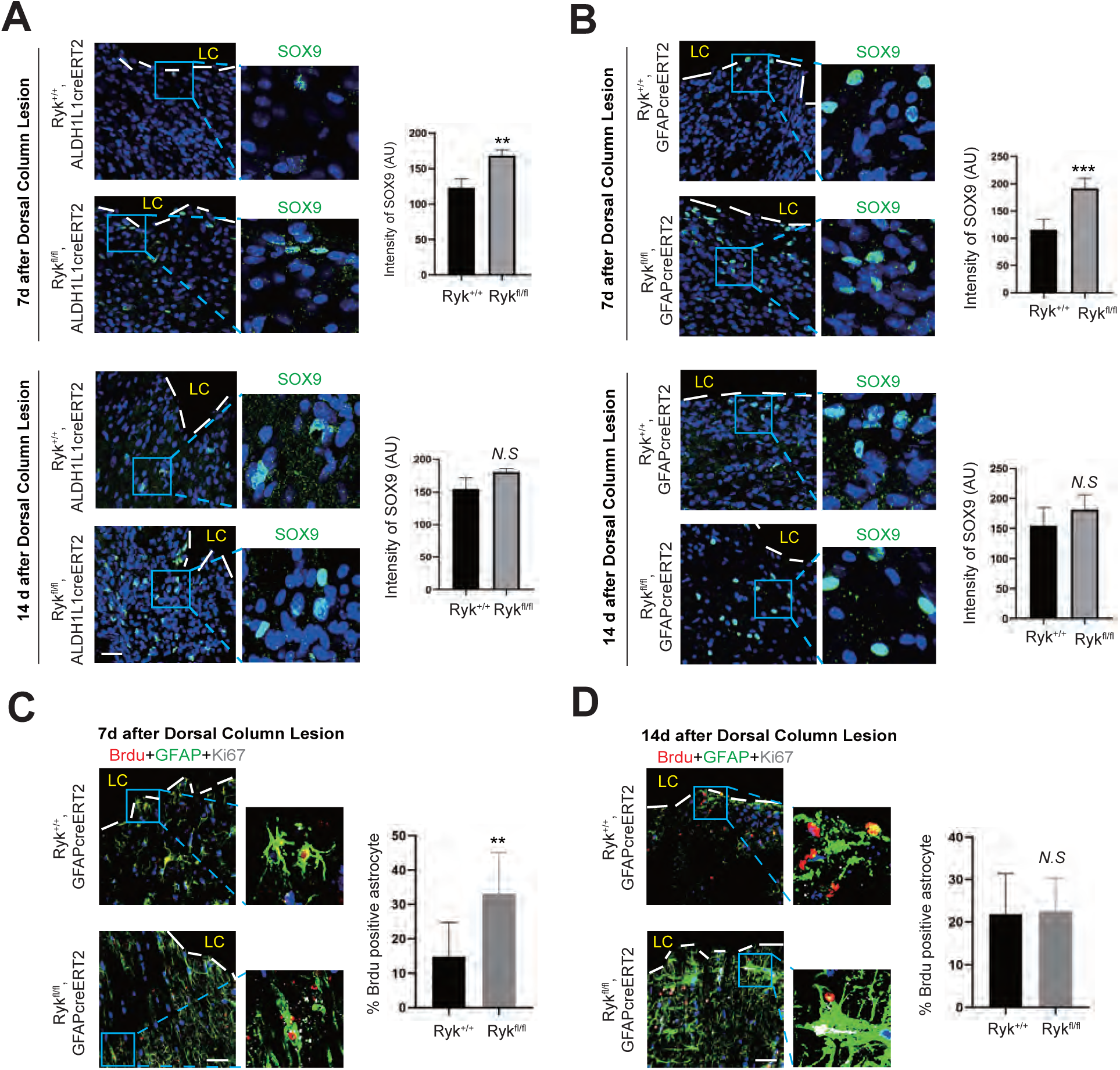
Accelerated astrocyte activation and proliferation around the lesion site in astrocyte-specific *Ryk cKO*. **(A)** Staining of spinal cord sections 7 days and 14 days after SCI with SOX9 antibody in *Ryk^fl/fl^* crossed with *Aldh1L1cre^ERT2^*. Scale bar =40 µm. Data are expressed as mean ± SD. N=3 for each group. \*\**P* < 0.01 and \*\*\**P* < 0.001 vs. control groups. **(B)** Staining of spinal cord sections 7 days and 14 days after SCI with SOX9 antibody in *Ryk^fl/fl^*crossed with *GFAPcre^ERT2^*. Scale bar =40 µm. Data are expressed as mean ± SD. N=3 for each group. \*\**P* < 0.01 and \*\*\**P* < 0.001 vs. control groups. **(C)** BrdU labeling and Ki67 staining in spinal cord sections 7 days after injury. Scale bar =40 µm. N=3 for each group. \*\**P* < 0.01 vs. control group. **(D)** BrdU labeling and Ki67 staining in spinal cord sections 14 days after injury. Scale bar =40 µm. N=3 for each group. \*\**P* < 0.01 vs. control group.

**Sup Fig 5.**
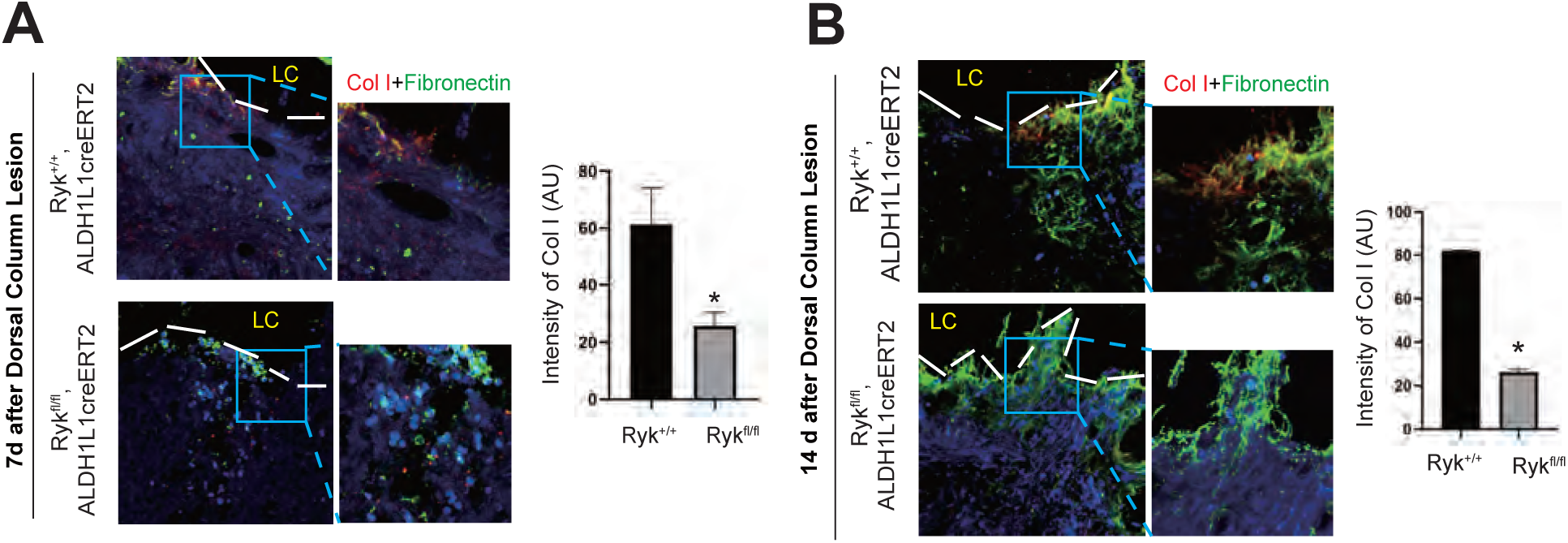
Changes of fibroblasts in astrocyte-specific *Ryk cKO*. (**A**) Decrease of immunoreactivity of Col I in astrocyte-specific *Ryk cKO* 7 days after injury. (**B**) Decrease of immunoreactivity of Col I in astrocyte-specific *Ryk cKO* 14 days after injury. Scale bar =40 µm. Data are expressed as mean ± SD. N=3 for each group. \**P* < 0.05 and \*\**P* < 0.05 vs. the indicated groups.

**Sup Fig 6.**
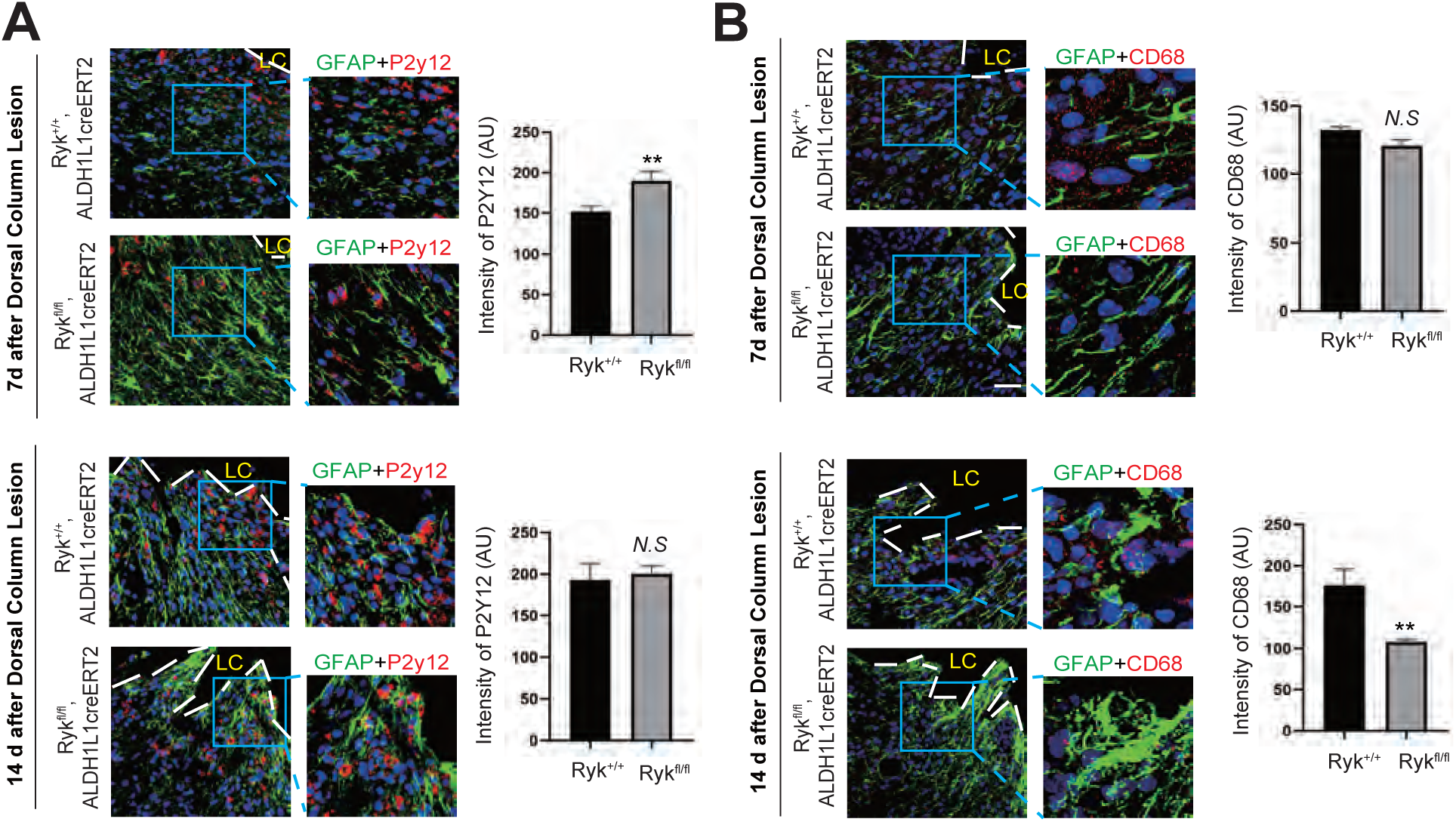
Changes of microglia in astrocyte-specific *Ryk cKO*. **(A)** Immunostaining of spinal cord sections with antibodies against GFAP and P2Y12 7 days or 14 days after C5 dorsal column lesion in control or astrocyte-specific *Ryk cKO*. **(B)** Immunostaining of spinal cord sections with antibodies against with antibodies against GFAP and CD68 7 days or 14 days after C5 dorsal column lesion in control or astrocyte-specific *Ryk cKO*. Scale bar =40 µm. N=3 for each group. Data are expressed as mean ± SD. \*\**P* < 0.01 vs. the indicated groups.

**Sup Fig 7.**
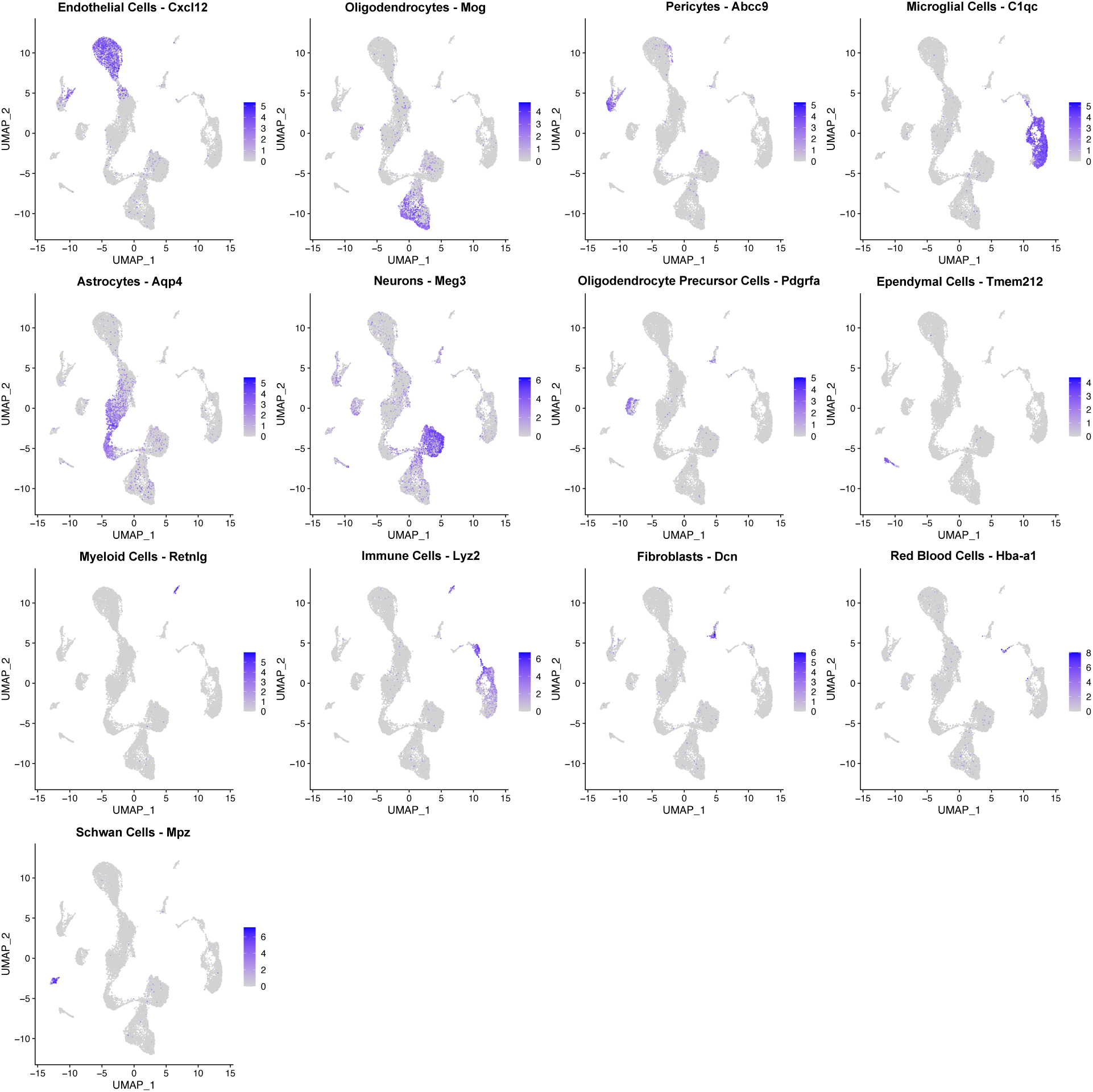
**Markers for major cell types.**

**Sup Fig 8.**
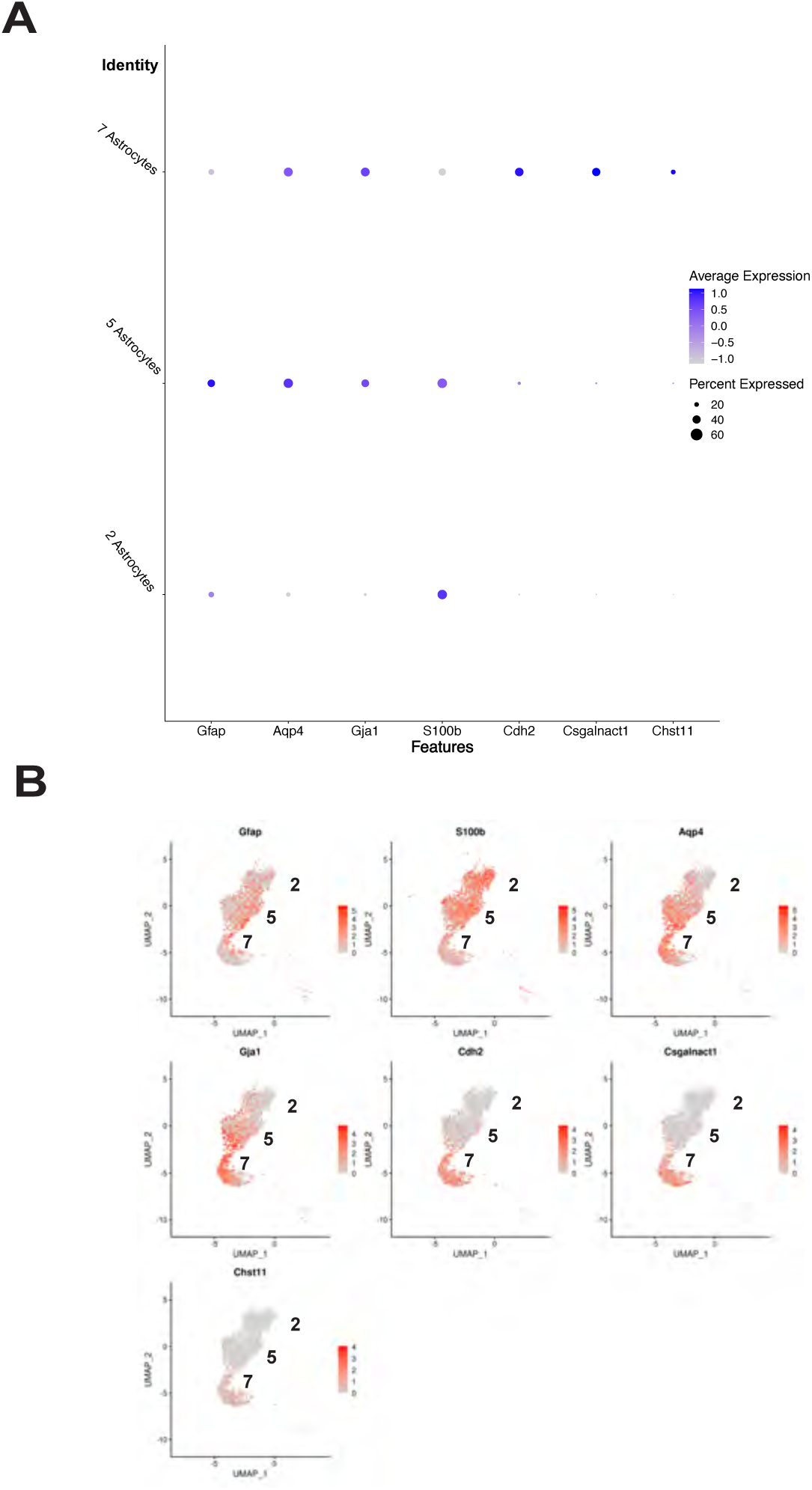
Makers for astrocyte subtypes. (**A**). Dot blots of markers in 3 subtypes of astrocytes. (**B**) Distribution of markers in 3 subtypes astrocytes.

**Sup Fig 9.**
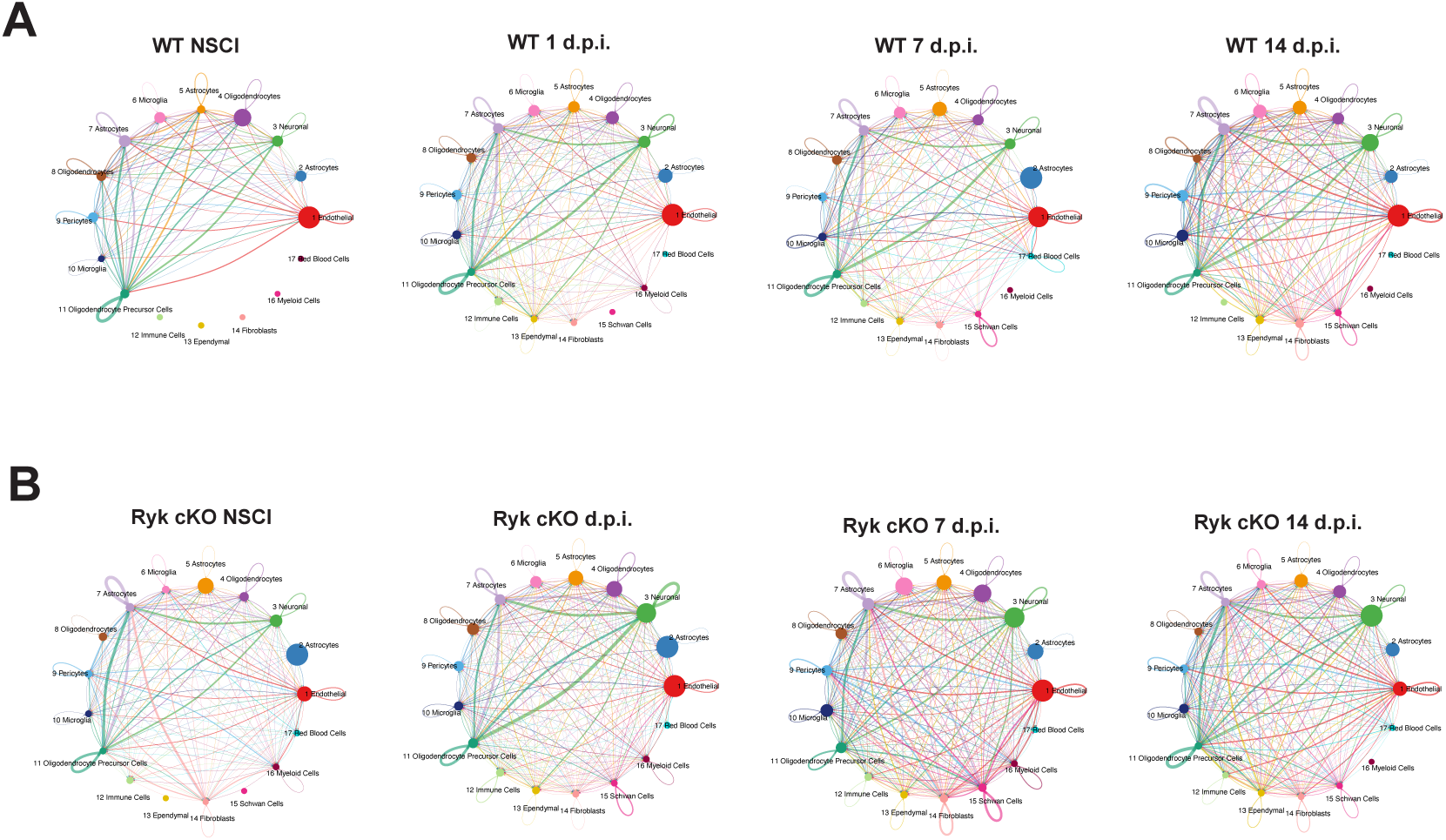
Changes of cell-cell communications after SCI. **(A)** Cell-cell communication network at different time points in control. **(B)** Cell-cell communication network at different time points in astrocyte-specific *Ryk cKO*.

**Sup Fig 10.**
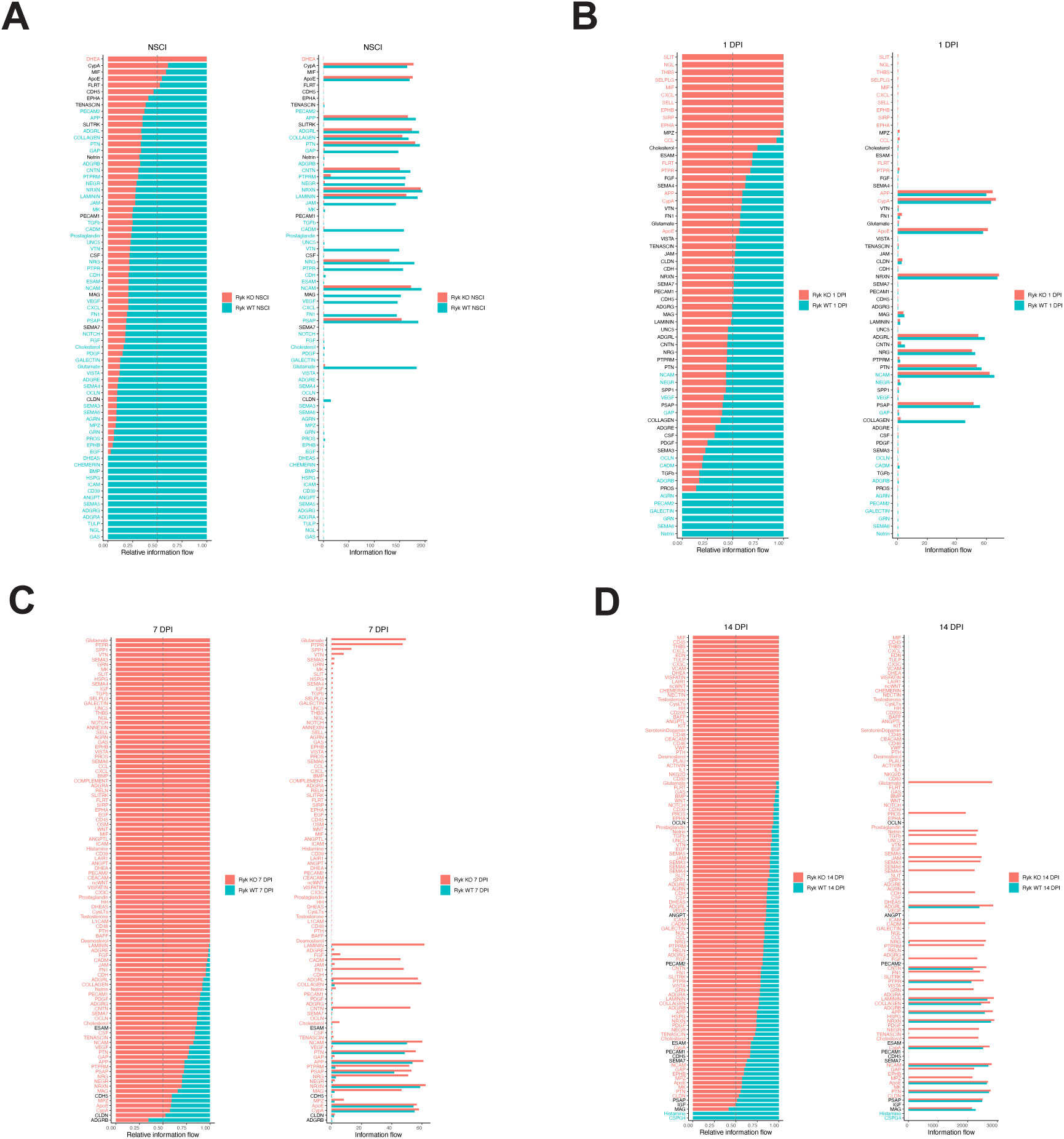
Ranking of signaling pathways which underwent the greatest changes in astrocyte-specific *Ryk cKO*. (**A**) Top pathways in astrocyte-specific *Ryk cKO* without spinal cord injury. (**B**) Top pathways in astrocyte-specific *Ryk cKO* 1 day post-injury. (**C**) Top pathways in astrocyte-specific *Ryk cKO* 7 days post-injury. (**D**) Top pathways in astrocyte-specific *Ryk cKO* 14 days post-injury.

**Sup Fig 11.**
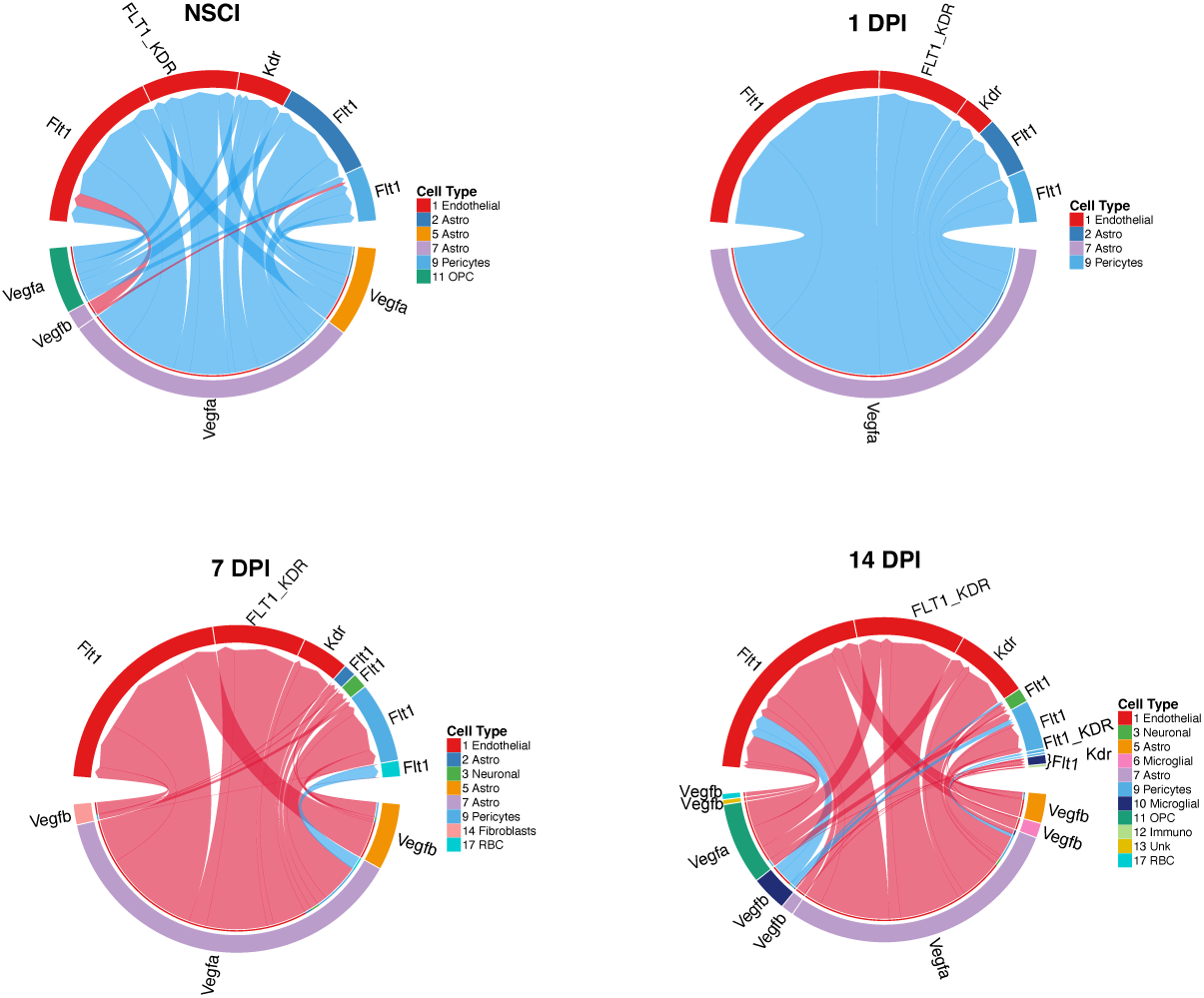
Changes of VGEF signaling in in astrocyte-specific *Ryk cKO*.

**Sup Fig 12.**
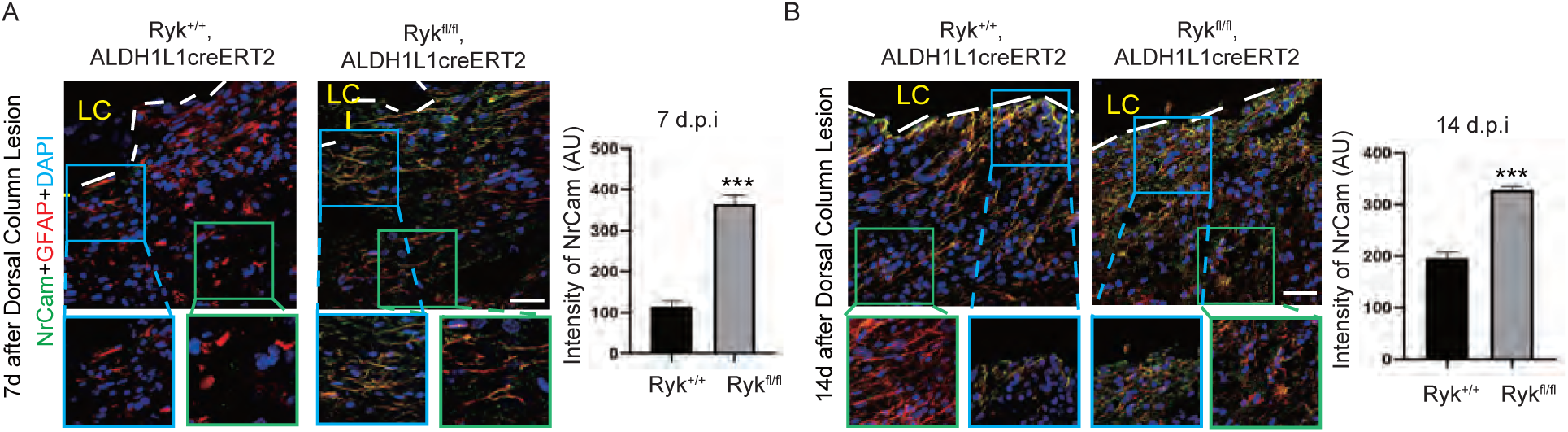
Changes of NrCAM expression in astrocyte-specific *Ryk cKO*. **(A)** Immunostaining of spinal cord sections with antibodies against NrCam and GFAP in control or *Ryk cKO* 7 days after injury. **(B)** Immunostaining of spinal cord sections with antibodies against NrCam and GFAP in control or *Ryk cKO* 14 days after injury.. Scale bar =40 µm. Blue box labelled the areas immediately abutting the lesion core while green box was about 100 µm away from the lesion core. Bar graphs showed quantitative analysis of the intensity of NrCam. N=3 for each group. Data are expressed as mean ± SD. \*\*\**P* < 0.001 vs. the indicated groups.

**Sup Fig 13.**
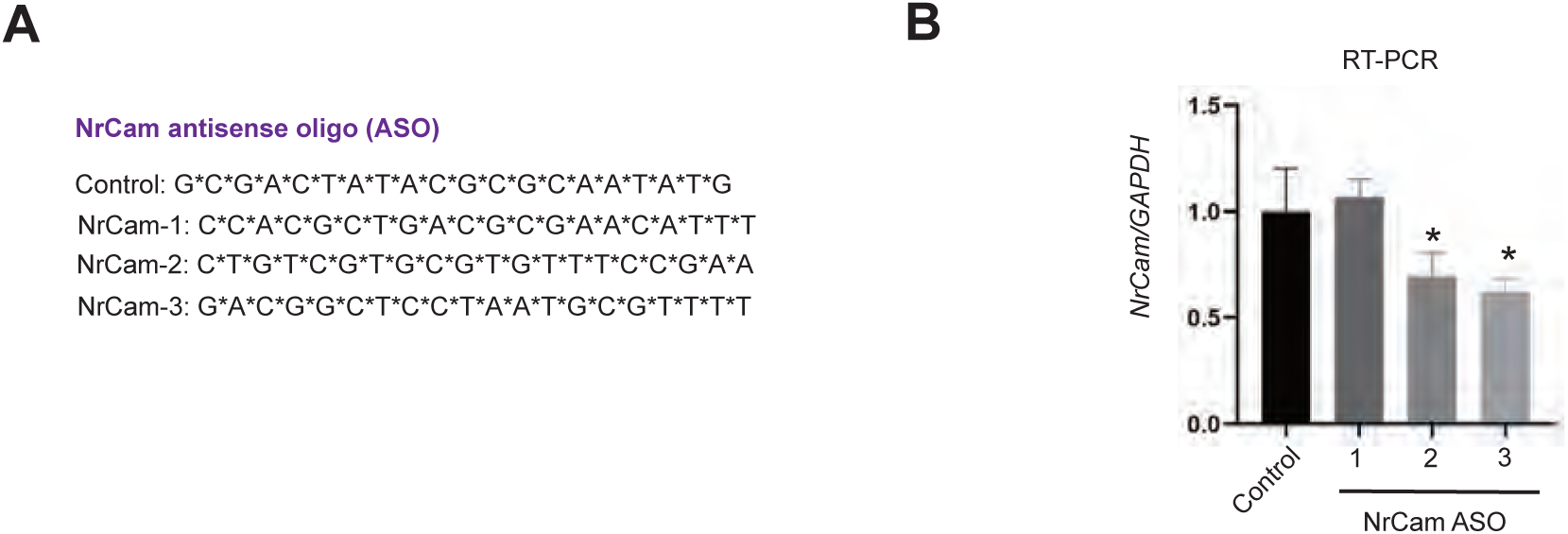
Design and validation of NrCAM antisense oligos. (**A**) Sequence of control and 3 ASOs against NrCAM. (**B**) RT-PCR to test the efficacy of antisense oligos.

